# Neural oscillation as a selective modulatory mechanism on decision confidence, speed and accuracy

**DOI:** 10.1101/2025.03.11.642591

**Authors:** Amin Azimi, KongFatt Wong-Lin

## Abstract

Neural oscillations have been associated with decision-making processes, but their underlying network mechanisms remain unclear. This study investigates how neural oscillations influence decision network models of competing cortical columns with varying intrinsic and emergent timescales. Our findings reveal that decision networks with faster excitatory than inhibitory synapses are more susceptible to oscillatory modulations. Higher in-phase oscillation amplitude reduces decision confidence without affecting accuracy, while decision speed increases. In contrast, anti-phase modulation increases decision accuracy, confidence and speed. Increasing oscillation frequency reverses these effects. Changing oscillatory phase difference gradually modulates decision behaviour, with decision confidence affected nonlinearly. Moreover, neural resonance can further enhance modulatory susceptibility for network with faster excitatory than inhibitory synapses. These effects decouple decision accuracy, speed and confidence, challenging standard speed-accuracy trade-off. These phenomena can be explained by excitatory neural populations contributing more to in-phase modulation, while inhibitory neural populations to anti-phase modulation. State-space trajectories’ momentum swinging with respect to network steady states and decision uncertainty manifold further provide insights into the neural circuit mechanisms. Our work provides mechanistic insights into how neurobiological diversity shapes decision-making processes in the presence of ubiquitous neural oscillations.

**Significance Statement:** Neural oscillations shape how the brain balances decision speed, accuracy, and confidence. Here, we show that tuning oscillatory amplitude, frequency, or phase can selectively alter these decision measures. In some cases, these modulations even break the usual trade-off between speed and accuracy, revealing a more flexible decision-making mechanism than previously assumed. Our findings highlight how synaptic time scales and rhythmic brain activity can give rise to distinct patterns of decision performance. This insight may guide new strategies for improving decisional processes in both healthy populations and clinical conditions.

## Introduction

Decision-making is a fundamental cognitive process involving the evaluation of options (Gold and Shadlen, 2007). A prevalent phenomenon of decision-making is the speed-accuracy trade-off (Fig. 1a), where faster decisions are made at the expense of decision accuracy, while more accurate decisions more likely take longer time to be made (Bogacz et al., 2010; Heitz, 2014). Further, decision confidence, a type of metacognition, generally tends to be higher with quicker decisions (Hellmann et al., 2024) or higher decision accuracy (Fig. 1a) (Jin et al., 2022).

**Fig. 1.**
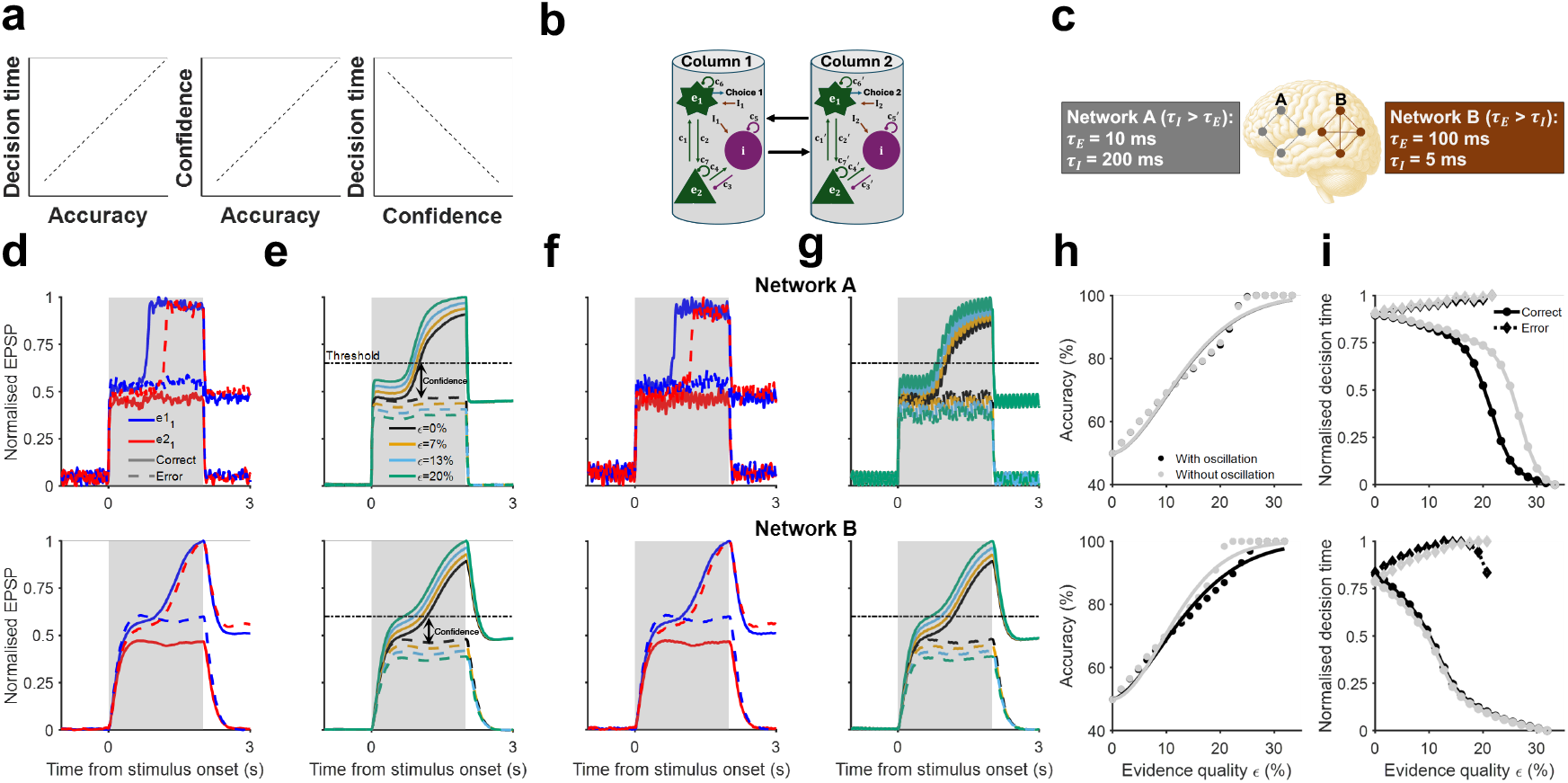
Models exhibit stereotypical choice behaviour in two-choice reaction time tasks. **a** Illustration of well-known trade-offs between decision accuracy, time and confidence. **b** Schematic of two-column cortical neural mass model for 2-choice decision-making. The choice is read out from population *e*_1_ in column 1 or column 2, corresponding to the random input. **c** Models incorporate neurophysiologically plausible synaptic time constants (network A with excitatory synaptic time constant (*τ*_*E*_) faster than that of inhibitory synapses (*τ*_*I*_); network B: *τ*_*E*_ > *τ*_*I*_). **d-i** Networks A (top) and B (bottom). **d** Sample single-trial simulation of normalised EPSP of neural population *e*_1_. Blue (red): correct (error) trials. Grey region: stimulus presence. *e*1_1_: EPSP of neuron *e*_1_ in column 1; *e*2_1_: EPSP of neuron *e*_1_ in column 2. **e** Normalised EPSP averaged over correct trials for different evidence qualities *ϵ*. Arrow: Decision confidence as the difference between output of winning column and losing column upon crossing decision threshold. **f** Simulated trials in the presence of oscillatory input (amplitude = 0.07 mA, frequency = 12 *Hz*). Solid (dotted): correct (error) trials. **g** Normalized EPSPs averaged over correct trials for different ϵ under oscillatory conditions. **h-i** Grey (black): with (without) oscillatory input. **h** Psychometric functions: decision accuracies vs. *ϵ*. **i** Chronometric functions: decision times vs. *ϵ*. Circle (diamond): correct (error) trials.

Decision making, and more generally, cognition, must be made amidst ongoing and ubiquitous neural oscillations—coherent fluctuations across frequency bands that coordinate neural activity (Fries, 2015). In fact, a key component of processing neuroimaging data in cognitive neuroscience involves the handling of such frequency bands to identify neural correlate for specific cognition (Buzsáki, 2006). In particular, different oscillatory frequency bands have been suggested to be associated with distinct functions. For example, beta-band oscillations (13 − 30 *Hz*) are linked to motor planning and decision-making (Engel and Fries, 2010; Baker et al., 2014) while gamma-band oscillations (30 − 100 *Hz*) to attention, working memory and decision-making (Pesaran et al., 2002; Siegel et al., 2012). Specifically, neural oscillations are crucial for integrating sensory input with motor actions, allowing adaptive cognitive state transitions (Wang, 2002, 2010). They shape decision-making by modulating neural communication timing and efficiency (Siegel et al., 2011), with phase synchronization enhancing information transmission speed and reliability (Buschman and Miller, 2007).

Non-invasive neuroimaging studies show that neural oscillations in gamma and beta bands predict perceptual decisions (Busch et al., 2009; Donner et al., 2009; Benwell et al., 2017). For example, (Wimmer et al., 2016) demonstrated that multiband oscillatory transitions in prefrontal circuits drive memory-guided decisions. Additionally, in visual discrimination tasks, alpha-band oscillations (8 − 13 *Hz*) modulate decision confidence without changing detection accuracy (Samaha et al., 2017, 2019). This implies that alpha oscillations might regulate the flow of sensory evidence during the decision-making process (Samaha et al., 2020).

Decision making often involves multiple brain regions and timescales (Soltani et al., 2021; Pinto et al., 2022). Moreover, sensory evidence accumulation timescale can vary across the cortex, which could be due to inherent differences in synaptic time constants (Wang et al., 2008) or network emergent phenomenon (Soltani et al., 2021) across the cortical hierarchy. Understanding how these differential dynamics modulate cognition in the presence of spontaneous neural oscillations is essential for comprehending their neural underpinnings in more realistic setting. Importantly, the computational neural circuit mechanisms of how neural oscillations influence decision-making are not well understood (Deco and Romo, 2008; Schmidt et al., 2018; Roach et al., 2023).

In this study, we address the above by first developing a cortical column based neural circuit model for decision making extended from previous models (Wong and Wang, 2006; Youssofzadeh et al., 2015). Our model comprises two excitatory and one inhibitory neural populations in each cortical column, with generated output encoding a potential decision outcome. When two cortical columns are coupled, the model can exhibit multistability that supports two-choice decision-making (Wilson and Cowan, 1972). We considered two networks with different sets of synaptic time constants: one with faster excitatory than inhibitory synapses and another with slower excitatory than inhibitory synapses. We simulate spontaneous neural oscillations as inputs to the neural circuit models. This approach allowed us to systematically evaluate how spontaneous network oscillations affect decision circuit endowed with different synaptic timescales. We first replicate experimental findings from relevant previous studies and then made various model predictions by varying the phases, amplitudes and frequencies of the neural oscillations. Our key findings are that neural oscillations can act as a novel and selective modulatory mechanism that can not only influence certain neural circuit type more but also can disentangle the tight relationship between decision speed, accuracy and confidence. Overall, our study highlights the intricate interplay between neural oscillations and intrinsic neurobiological timescales in modulating decision dynamics.

## Materials and Methods

### Cortical columnar neural mass model for decision-making

We develop a two-column cortical model for two-alternative decision-making based on our fully self-feedback neural mass model (Youssofzadeh et al., 2015), The model comprises two excitatory neural populations (*e*_1_ and *e*_2_) and one inhibitory neural population (i) in each column and these neural populations are rate-based. In line with biologically realistic cortical column architectures for modelling neuroimaging data (David et al., 2006; Youssofzadeh et al., 2015), our model incorporates two excitatory neural populations—spiny stellate cells and pyramidal cells—together with one inhibitory neural population—inhibitory interneurons. Although for simplicity, the two excitatory populations (*e*_1_ and *e*_2_) share identical neuronal properties (see Supplementary Fig. S1), their explicit inclusion reflects the cellular diversity observed in cortical layers and they have different afferent and efferent connectivity. This design provides a biologically grounded basis for the model and distinguishes it from simplified approaches that use only a single excitatory population (e.g., Roach et al., 2023; Najafi et al., 2020).

This model integrates inputs into both the excitatory and inhibitory populations, and incorporates lateral interactions between the columns, as depicted in Fig. 1b (see Fig. S1 for details), allowing winner-take-all behaviour that is required for forming unitary decision.

### Postsynaptic potential function

In this model, the function *S*, characterized by a sigmoidal shape, represents the postsynaptic potentials and is defined as:

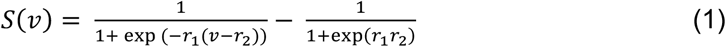

where *r*_1_ (= 2) and *r*_2_ (= 1) are parameters controlling the characteristics of the sigmoid function; increasing *r*_1_ steepens the slope, while increasing *r*_2_ shifts the curve to the right.

The average membrane depolarisation *v* (or postsynaptic potentials) associated with a neural population serves as the presynaptic input for other connected neural populations, including other cortical columns, through the connections as described by *S*(*v*) (Youssofzadeh et al., 2015). The priors and physiological interpretations of the model parameters are provided in Table 1.

**Table 1:** Priors and physiological interpretations of model parameters. Subscript *e* (*i*) denote excitatory (inhibitory). Compare with Fig. S1.

| Parameter | Physiological interpretation | Value |
| --- | --- | --- |
| $H_e, H_i$ | Excitatory and inhibitory magnitude in synaptic function | 3.25, 22 [mV] |
| $K_e = \frac{1}{\tau_e}, K_i = \frac{1}{\tau_i}$ (Network A) | Excitatory and inhibitory rate constant (AMPA, GABA<sub>B</sub>) | 100, 2 [$s^{-1}$] |
| $K_e = \frac{1}{\tau_e}, K_i = \frac{1}{\tau_i}$ (Network B) | Excitatory and inhibitory rate constant (NMDA, GABA<sub>A</sub>) | 10, 100 [$s^{-1}$] |
| $K_e = \frac{1}{\tau_e}, K_i = \frac{1}{\tau_i}$ (Network C) | Excitatory and inhibitory dynamical rate constant | 50, 100 [$s^{-1}$] |
| $c_1, c_2, c_3, c_4$ (Network A) | Intrinsic coupling parameters for each column | 0.1, 0.1, 8, 5 |
| $c_1, c_2, c_3, c_4$ (Network B) | Intrinsic coupling parameters for each column | 0.15, 0.1, 32, 55 |
| $c_1, c_2, c_3, c_4$ (Network C) | Intrinsic coupling parameters for each column | 0.1, 0.1, 8, 5 |
| $c_5, c_6, c_7$ (Network A) | Intrinsic coupling parameters in feedback loops for each column | 0.3, 1, 105 |
| $c_5, c_6, c_7$ (Network B) | Intrinsic coupling parameters in feedback loops for each column | 50, 0.1, 31.5 |
| $c_5, c_6, c_7$ (Network C) | Intrinsic coupling parameters in feedback loops for each column | 0.3, 1, 80 |
| $c_{ee}, c_{ei}, c_{ee}', c_{ei}'$ (Network A) | Extrinsic coupling parameters (lateral connections) | 0.7, 30, 0.7, 30 |
| $c_{ee}, c_{ei}, c_{ee}', c_{ei}'$ (Network B) | Extrinsic coupling parameters (lateral connections) | 0.85, 50, 0.85, 50 |
| $c_{ee}, c_{ei}, c_{ee}', c_{ei}'$ (Network C) | Extrinsic coupling parameters (lateral connections) | 0.7, 30, 0.7, 30 |

### Model parameters

### Model equations

Fig. 1b (see Fig. S1 for details) illustrates that neural population *e*_2_ in the first column is laterally connected to neural population *e*_1_ in the second column, and vice versa. Additionally, there are lateral connections between neural population *e*_2_ in the first column and neural population *i* in the second column, and vice versa. Accordingly, the dynamical equations describing the model can be described as follows (see Table 1 for notation meaning).

For cortical column j (the other column denoted as k):

Post-synaptic potential for *e*1_*j*_:

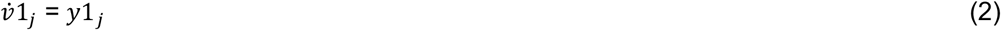

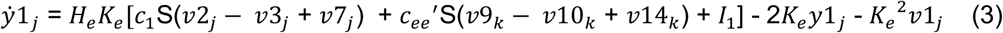

Post-synaptic potential for *e*2_*j*_:

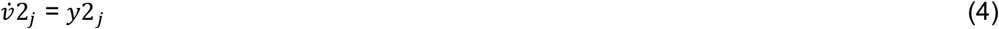

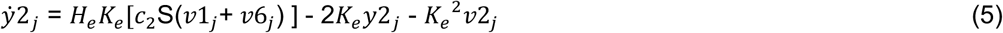

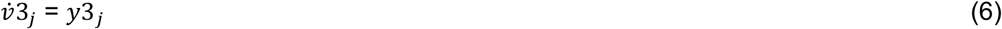

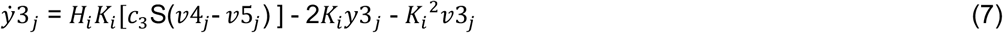

Post-synaptic potential for *i*_*j*_:

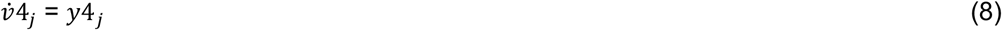

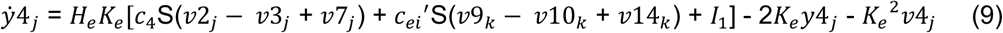

Post-synaptic potential due to self-inhibition for *i*_*j*_:

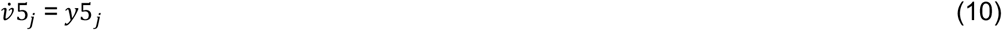

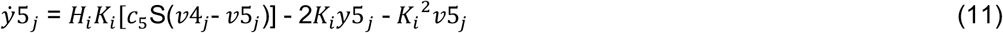

Post-synaptic potential due to self-excitation for *e*1_*j*_:

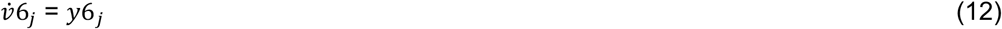

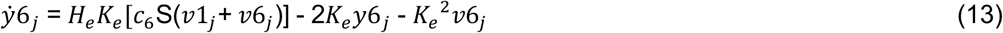

Post-synaptic potential due to self-excitation for *e*2_*j*_:

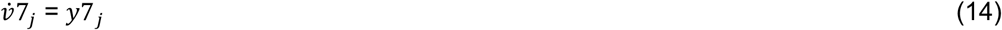

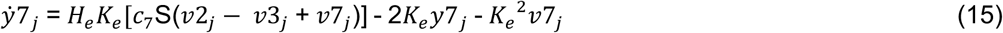

### Model input

Then, we model the stimulus input to both columns using a Gaussian bump function (Friston, 2008):

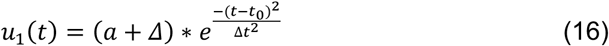

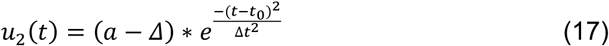

Here, *a* represents the amplitude for zero bias (Δ), *t*_0_=1 s is the stimulus onset time, t=3 s is the stimulus offset time, and *Δ*t (0.007 s) controls the width of the Gaussian bump. Additionally, we measure the evidence quality ε (analogous to motion coherence (Atiya et al., 2019)) using the following definition:

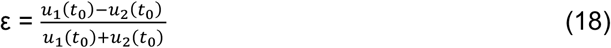

Also, noise is incorporated into the dynamics of the excitatory *e*_1_ and inhibitory *i* populations via inputs *I*_*η*_, processed through rapid synaptic activation mediated by AMPA receptors. The dynamics governing this interaction is described by (Destexhe et al., 2001):

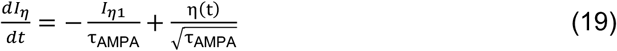

Here, τ_AMPA_= 5 ms, and *I*_*η*_ is a white Gaussian noise with zero mean and standard deviation of 0.02 nA.

In addition, background input (*I*_0_) and sinusoidal stimulation (*I*_*O*_) at a frequency f (12 *Hz*) are incorporated into the background noise of the inputs received by both cortical columns, mimicking spontaneous brain oscillations (Buzsáki and Draguhn, 2004). The total input current to each of the columns is therefore:

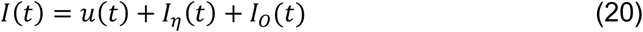

Unless specified, this total input is applied with equal strength to both excitatory and inhibitory units within each column.

### Model analysis

We numerically integrate the nonlinear equations of the model using the Euler-Maruyama method in MATLAB R2023b, with a temporal resolution of 1 ms. Simulations are conducted to investigate two-alternative decision-making and decision storage through multistable dynamics. We manipulate the intrinsic and lateral inhibitory and excitatory connections to examine the multistability of the postsynaptic potentials in *e*_1_ as influenced by input currents, utilising XPPAUT 8.0 for these simulations (Ermentrout and Mahajan, 2003).

Due to the symmetry of the cortical columns, we assume without loss of generality that activity in the first column encodes the correct choice. Specifically, we increase the input to the first column relative to the second column. We then classify the simulated trials into three categories: (1) correct trials, characterised by a significantly higher output from the first column; (2) error trials, identified by a higher output from the second column; and (3) non-decision trials, defined by an output difference that do not surpass a predefined threshold (network A: 0.01 mV; network B: 0.08 mV).

In the simulations for both networks, an evidence quality of 7% was used, and 3000 trials were generated for each frequency–amplitude condition tested. Without oscillatory input, Network A generated 1807 correct trials, 1063 error trials, and 130 non-decision trials, whereas Network B generated 1815 correct trials, 985 error trials, and 200 non-decision trials. For both networks, the simulation data were not mean-centered. Also, each decision parameter (accuracy, decision time, and decision confidence) was computed across 21 amplitude and frequency values under different conditions.

### Phase-locking analysis

In the phase-locking analysis, the instantaneous phase of the output signal—band-pass filtered in the alpha frequency range (8–12 *Hz*)—was estimated using the Hilbert transform to determine the phase difference between output peaks and the corresponding epochs of the input signal (Azimi et al., 2021).

### Bootstrapped Confidence Intervals (CI)

Confidence intervals were estimated using a nonparametric bootstrap procedure with 1000 resamples of simulated outcomes. For each resample, decision measures (accuracy, decision time, decision confidence) were recalculated, and the 2.5th and 97.5th percentiles were taken as bounds of the 95% confidence interval.

### Statistical Analysis

Statistical analyses are performed using MATLAB R2023b. All statistical tests are calculated using Pearson correlation coefficient, with statistical significance set at *p < 0*.*05*.

For the calculation of partial correlations between oscillatory input amplitude and decision parameters, different frequency bins were considered, and within each bin the Pearson correlation and corresponding p-values between input amplitude and decision parameters were reported. A similar procedure was applied for calculating partial correlations between oscillatory input frequency and decision parameters.

### Data and Code Availability

The simulations and figure generation for this study are conducted using MATLAB, and all the associated code is openly available in our GitHub repository at https://github.com/amin8179/Impact-of-oscilllation-on-decision-making/tree/main. The repository includes a detailed “README” file that provides comprehensive instructions on how to reproduce all the figures presented in the main manuscript and the Supplementary Information. Since all data are generated through simulations of the model, no empirical data collection is necessary. The “README” file also outlines clear steps for generating the simulation data shown in this study.

## Results

### Decision networks operating at different timescales

Two-column cortical model for two-alternative decision-making endowed with neurophysiologically informed synaptic timescales and multistable dynamics was developed, building on our previous fully self-feedback neural mass model (Youssofzadeh et al., 2015). Each column comprised two excitatory neural populations and one inhibitory neural population (Fig. 1b; see Fig. S1a for details), each with self-feedback connectivity.

Additionally, in our model, stimulus inputs were applied to both the inhibitory population (*i*) and an excitatory (*e*_1_) population in each column. While Najafi et al., 2020 demonstrated that inhibitory neurons can be decision-selective, it remains unclear whether selective sensory inputs directly target both excitatory and inhibitory populations. To reflect broader possibilities during sensory evidence integration, we therefore implemented selective sensory input to both excitatory and inhibitory neural populations within cortical columns, rather than exclusively to excitatory neural populations. In later analyses, we further disentangled their contributions by selectively applying inputs to either excitatory or inhibitory populations alone. Due to cross-columnar inhibition (Figs. 1b and S1a), the model exhibited winner-take-all behaviour between the competing columnar neural population activities. Further, self-feedback allowed persistent activities for storage of decision made over time.

Inspired by the cortical column models of David et al., 2006 and Youssofzadeh et al., 2015, which have been used for brain-wide modelling, our implementation features a biologically realistic architecture with two classes of excitatory neural populations —spiny stellate cells and pyramidal cells—and a single population of inhibitory interneurons, thereby capturing the cellular diversity characteristic of interacting cortical layers.

We further developed the above model into two types, A and B, each with different timescales characterised by different sets of synaptic time constants (Gerstner et al., 2014). Specifically, network A was presumed to be dominated by faster (e.g. AMPA-mediated) excitatory than (e.g. GABA_B_-mediated) inhibitory synapses, while network B was dominated by slower (e.g. NMDA-mediated) excitatory than inhibitory (e.g. GABA_A_-mediated) synapses (Fig. 1c) (see Methods). The values of the synaptic time constants were selected based on known physiological ranges (Bal and Destexhe, 2009).

Upon stimulus presentation, both networks can produce distinct unitary choice outcomes with increasing activity for one of the excitatory neural populations (Figs. 1b and S1a, populations *e*_1_) in either competing cortical columns. Specifically, these circuits indicate a choice by having a columnar neural population*’*s excitatory postsynaptic potential (EPSP) to ramp up over time towards a prescribed decision threshold, akin to temporal integration of evidence prior to choice commitment (Fig. 1d). The decision time was defined as the time from stimulus onset to decision threshold crossing of the “winning” neural population, and decision speed is inversely linked to this time. The confidence associated with the decision was computed at the decision time based on the difference between the activities of the winning column and the losing column which did not cross the threshold (Kepecs et al., 2008; Vugt et al., 2014) (Fig. 1e). Further, the EPSP of the winning excitatory population increased more rapidly with higher evidence quality *ε* (indicating easiness level of task, e.g. equivalent to motion coherence in standard random-dot kinetogram (Atiya et al., 2019); see Methods) (Fig. 1e).

Motivated by a previous experimental study^21^ which has identified the relationship between alpha oscillation (8 − 13 *Hz*) with (perceptual) decision confidence, we first applied oscillatory input of 12 *Hz* (0.07 *mA*) on both networks A and B. We found that introducing 12 *Hz* neural oscillations preserved the core decision-making and working memory behaviours of the network models (Fig. 1f, g). Simulated trials in which a clear separation between the columnar EPSPs (difference of at least 0.01 *mV* for network A; 0.08 *mV* for network B) failed to occur are deemed invalid and excluded from consideration for both the psychometric and chronometric functions; this occurred only for about 5% for networks A and B in the study.

As evidence quality increased, the models’ choice accuracy monotonically increased from chance level (50%) towards 100% (Fig. 1h; black line), while decision time decreased, with slower incorrect than correct choices (Fig.1i; black line), in agreement with stereotypical choice behaviour (Shadlen and Newsome, 2001; Gold and Shadlen, 2007). Additionally, persistent EPSP activity of the winning neural population after stimulus offset allowed for choice readout even after significant time delay by invoking working memory to store the decision made, consistent with previous studies (Shadlen and Newsome, 2001; Roitman and Shadlen, 2002; Wang, 2002; Wong and Wang, 2006; Roach et al., 2023). Notably, our study demonstrates that network with faster excitatory than inhibitory synapses could still exhibit decision-making behaviour, challenging previous neurobiologically realistic computational modelling studies (Wang, 2002; Wong and Wang, 2006; Roach et al., 2023). Interestingly, the initial rise time of such network’s competing activities was rapid prior to their splitting (Friston, 2008). Furthermore, applying 12 *Hz* neural oscillations could alter the psychometric and chronometric functions (Fig. 1h, i; grey line), with decision accuracy being more resilient to neural oscillatory modulation. This hinted that neural oscillation might potentially disentangle decision’s speed–accuracy trade-off (Bogacz et al., 2010; Heitz, 2014), which we shall demonstrate with further evidence below. Moreover, neural population *e*_2_ shares the same dynamical properties as population *e*_1_ in both networks, as evidenced by the bistable decision-making dynamics shown in Supplementary Figures S1b.

### Distinct oscillatory entrainment for decision networks with different timescales

As described above, we applied a 12 Hz sinusoidal drive simultaneously to the principal excitatory neural population (*e*_1_) and the inhibitory neural population (*i*) in two distinct network architectures (Networks A and B; Figs. S2a and e) to assess how rhythmic inputs shape local network dynamics. In Network A, *e*_1_ exhibited robust entrainment: the membrane-potential trace in column 1 closely followed the sinusoidal drive, producing large-amplitude oscillations phase-locked to the input (Fig. S2a, bottom). Spectral analysis confirmed this observation, as both the injected drive and the *e*_1_ output displayed a prominent alpha-band peak at ∼12 Hz in their power-spectral densities (PSDs; Fig. S2b). Quantitatively, cross-correlation analysis between the drive and the *e*_1_ output revealed strong coupling, with the peak correlation occurring at a lag of ∼25 ms (Fig. S2c), indicating a consistent temporal relationship. Moreover, phase-locking analysis yielded a mean resultant vector length of 0.9927 (Rayleigh test), with a preferred phase of 84.1°, demonstrating nearly perfect locking of the oscillatory amplitude to a specific phase of the input rhythm (Fig. S2d).

In contrast, Network B exhibited markedly weaker entrainment to the same 12 Hz drive. The *e*_1_ membrane-potential trace showed only faint modulation by the input (Fig. S2e, bottom). Although the drive’s PSD retained a clear ∼12 Hz peak, the PSD of the e_1_ output shifted to ∼14 Hz and displayed reduced power at 12 Hz (Fig. S2f). Cross-correlation analysis further revealed a pronounced decrease in input–output coherence (Fig. S2g). Finally, time–frequency decomposition of the *e*_1_ output uncovered only weak, intermittent alpha-band activity throughout the simulation (Fig. S2h). Together, these results demonstrate that—even under identical rhythmic stimulation—Networks A and B differ fundamentally in their capacity to track and reshape oscillatory inputs.

### In-phase oscillation amplitude distinctly affects decision networks with different timescales

As previous experimental studies have shown that neural oscillations in the alpha frequency band (8 − 13 *Hz*) could influence perceptual decisions (Samaha et al., 2017), we first applied in-phase sinusoidal modulation at a frequency of 12 *Hz* to both cortical columns in both networks, simulating influence from spontaneous neural oscillations (Fig. 2a, *ϵ* = 7%). To compare the changes in decision time and confidence after modulation of oscillatory input, their values were normalised to their maximum values. For network A which had faster excitatory than inhibitory synapses, the 12 *Hz* in-phase oscillatory modulation did not affect decision accuracy (Fig. 2b, dark grey) but decision confidence could be reduced by as much as about 15% (Fig. 2d, dark grey) for correct (*r* = −0.89, *p* = 3.1 × 10^−8^, 95% *CI* [0.945, 0.983]) and error trials (*r* = −0.86, *p* = 4.2 × 10^−7^, 95% *CI* [0.991, 0.998]). This was consistent with experimental findings that associate alpha oscillations with decreased confidence (Samaha et al., 2017, 2020) in both correct and error trials while not affecting decision accuracy (Samaha et al., 2017, 2020). Additionally, decision time was predicted to be shortened by as much as about 5% for correct (*r* = −0.90, *p* = 2.1 × 10^−8^, 95% *CI* [0.989, 0.996]) trials (Fig. 2c, dark grey) and it shows no significant change for error trials.

**Fig. 2.**
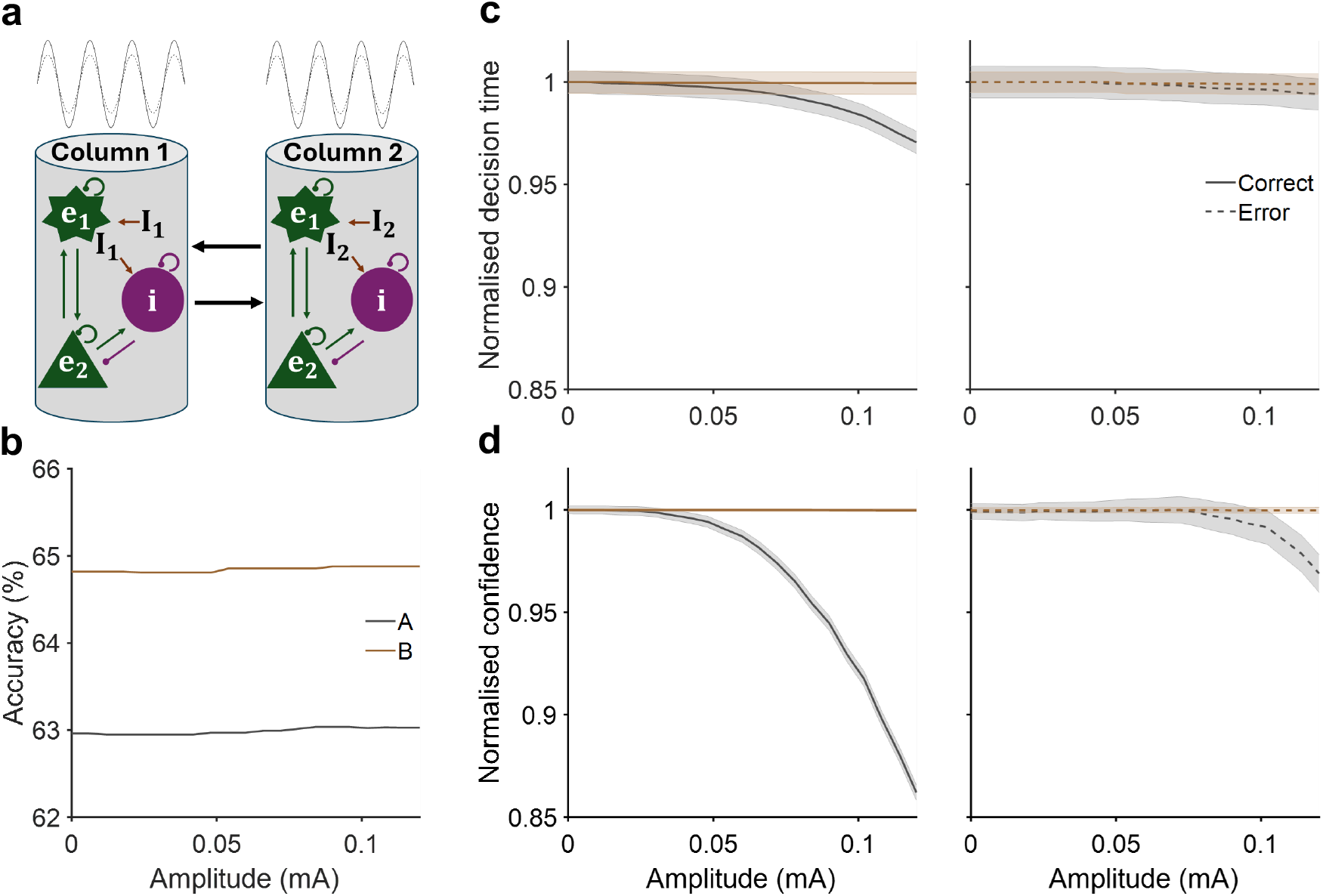
Different effects of in-phase 12 *Hz* neural oscillatory amplitude on networks with different timescales. **a** In-phase sinusoidal modulation at different amplitudes applied to both cortical columns for each network. **b** Decision accuracy remained unchanged for both networks. **c** In Network A (grey), increasing oscillation amplitude shortened normalised decision time for correct trials but produced no significant change for error trials; Network B (brown) showed no significant change for either trial type. **d** For network A, normalised decision confidence decreased with higher oscillation amplitudes; network B showed no significant changes for both correct (left) and error (right) trials.

For network B which had slower excitatory than inhibitory synapses, decision accuracy remained unchanged across varying oscillation amplitudes (Fig. 2b, brown), indicating that in-phase oscillatory activity did not affect the correctness of decisions made by the network. Additionally, increasing the oscillation amplitude did not significantly affect the decision times for either correct or error trials (Fig. 2c; brown). Moreover, there were no considerable changes in decision confidence observed in network B for both correct and error trials across different oscillation amplitudes (Fig. 2d; brown). This indicated that, at 12 *Hz* oscillation frequency, increasing the oscillation amplitude did not affect decision accuracy, time, or confidence for network B.

### In-phase oscillation frequency effects are opposite that of amplitude modulation

Next, we investigate the effects of neural oscillation frequency on decision-making behaviour by gradually increasing the input frequency to both networks A and B (Fig. 3a, amplitude=0.08 *mA*). Our results indicated that higher frequencies did not noticeably alter decision accuracy in either network (Fig. 3b). Further, increasing the frequency did not significantly alter either decision time or confidence in network B across both trial types (Fig. 3c–d). In contrast, in network A (Fig. 3c–d), decision confidence increased by approximately 5% (*r* = 0.69, *p* = 1.1 × 10^−3^, 95% *CI* [0.986, 0.997]) and decision time by about 1% (*r* = 0.75, *p* = 2.1 × 10^−4^, 95% *CI* [0.996, 0.999]) for correct trials, with no significant changes observed for error trials. These findings demonstrate that the alpha to gamma range for Network A more effectively influence both decision time and confidence while maintaining decision accuracy.

**Fig. 3.**
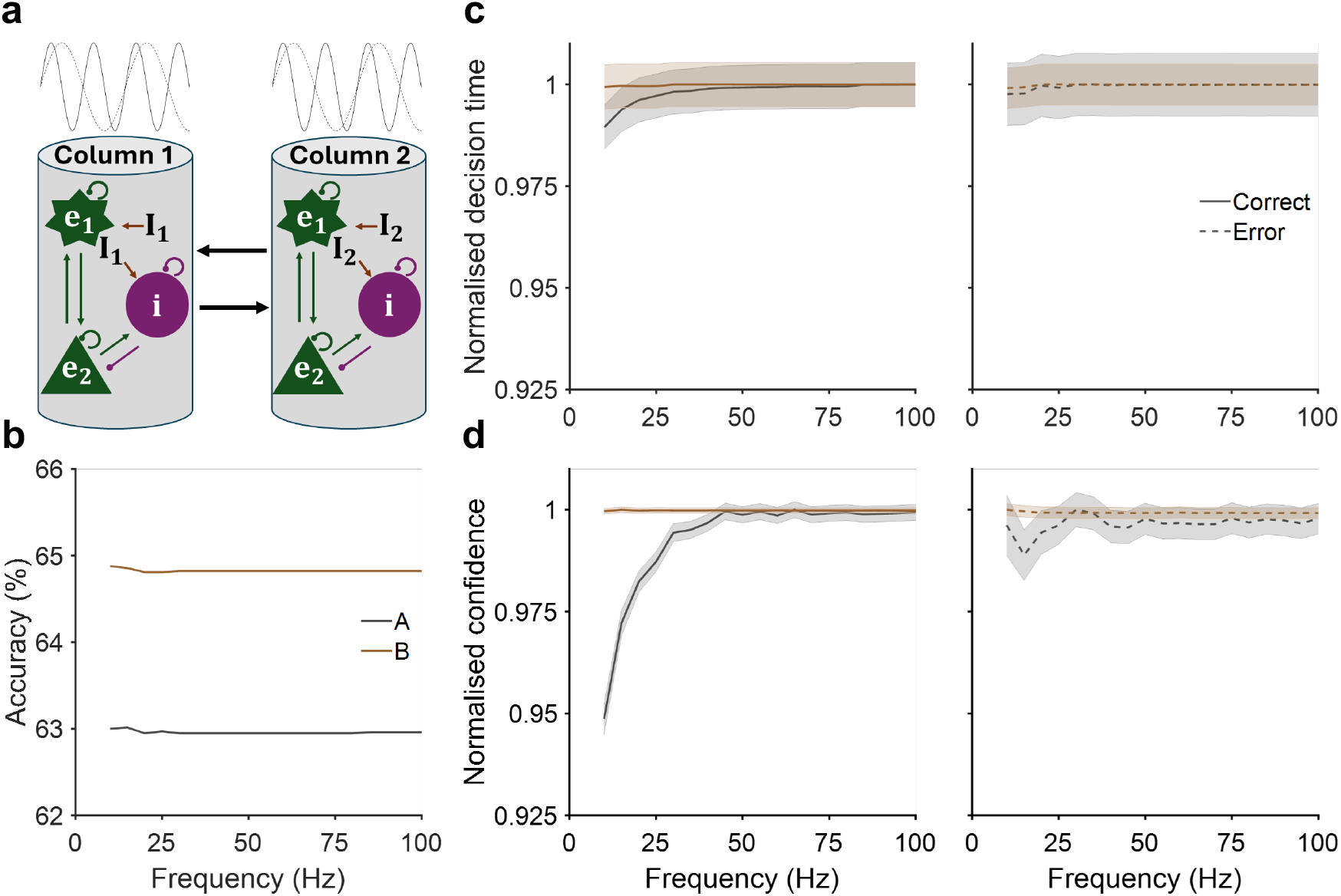
Different effects of in-phase neural oscillatory frequency (amplitude = 0.08 *mA*) on networks with different timescales. **a** In-phase sinusoidal modulation at different frequencies applied to both cortical columns for each network. **b** Decision accuracy remained unchanged for both networks. **c, d** For Network A, normalized decision time (c, grey) and confidence (d) increased for correct trials (left) but showed no substantial change for error trials (right). In contrast, for Network B, neither decision time nor confidence changed significantly for either type of trial (c–d, brown).

### Anti-phase oscillation amplitude enhances decision efficiency

Neural oscillation may not necessarily have to affect different brain locations synchronously. For instance, this may be due to transduction or conduction delays in receiving neural signals (Sanchez Bornot et al., 2018) (see Discussion for further potential mechanisms). Hence, we next examine the other extreme case of neural oscillations with anti-phase sinusoidal modulation between the two competing cortical columns. We began by applying oscillatory modulation at a frequency of 12 *Hz* (alpha band) to both cortical columns within each network (Fig. 4a, grey). For network A, enhancing the amplitude of anti-phase oscillatory modulation led to improved decision accuracy (by ∼40%) (Fig. 4b, dark grey, *r* = 0.98, *p* = 1.2 × 10^−15^, 95% *CI* [0.783, 0.875]), decreased decision time (by ∼40%) for correct trials (*r* = −0.99, *p* = 1.0 × 10^−21^, 95% *CI* [0.853, 0.918]), and slight increase of decision time (by ∼10%) for error trials (Fig. 4c, dark grey, *r* = 0.84, *p* = 1.8 × 10^−6^, 95% *CI* [0.937, 0.963]). Additionally, decision confidence increased for both trial types (Fig. 4d, dark grey, *r* = 0.92, *p* = 2.2 × 10^−9^, 95% *CI* [0.873, 0.913] for correct, *r* = 0.64, *p* = 2.0 × 10^−3^, 95% *CI* [0.946, 0.960] for error). This suggests that anti-phase oscillations can facilitate more accurate and faster decisions, unlike standard speed-accuracy trade-off mechanisms.

**Fig. 4.**
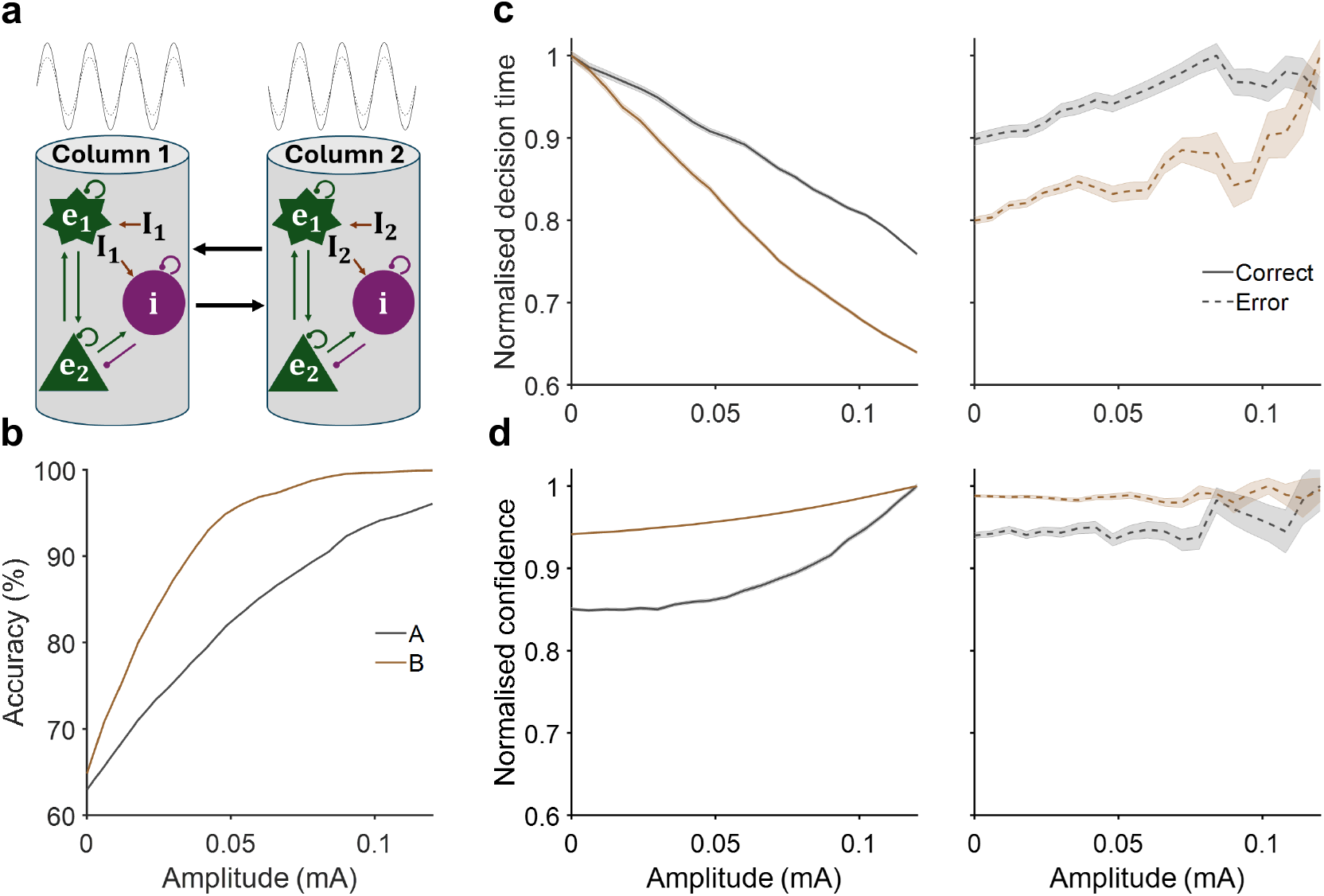
Anti-phase 12 *Hz* neural oscillatory amplitude enhances decision accuracy and modulates decision speed. **a** Anti-phase sinusoidal modulation with varying amplitudes applied to both cortical columns for each network. **b** Decision accuracy improved with higher oscillation amplitudes in both networks. **C** Normalised decision time decreased with increasing amplitude for correct trials (left), while increasing for error trials (right). **d** Normalized decision confidence increased in Network A for both correct and error trials, as well as for correct trials in Network B. However, no substantial change was observed in Network B for error trials.

For network B, increasing the amplitude of anti-phase oscillatory modulation resulted in enhanced decision accuracy (by ∼40%) (Fig. 4b, brown, *r* = 0.88, *p* = 1.5 × 10^−7^, 95% *CI* [0.872, 0.957]), decreased decision time for correct trials (by ∼40%) (*r* = −0.99, *p* = 1.0 × 10^−21^, 95% *CI* [0.755, 0.854]) and increased decision time for error trials (by ∼20%) (*r* = 0.85, *p* = 1.4 × 10^−6^, 95% *CI* [0.842, 0.882]), and increased decision confidence for correct trials (*r* = 0.98, *p* = 1.8 × 10^−15^, 95% *CI* [0.957, 0.973]), while maintaining stable decision confidence across error trials. The slower error decisions led to further temporal accumulation of evidence and enhancing evidence quality. Thus, this suggests that anti-phase neural oscillations can make more efficient decisions regardless of the network’s timescale.

### Anti-phase oscillation frequency effects are opposite that of amplitude modulation

As in the in-phase oscillation modulation investigation, here we increased the frequency of anti-phase oscillatory modulation and observed longer decision time for correct trials (amplitude = 0.08 *mA, r* = 0.80, *p* = 4.0 × 10^−5^, 95% *CI* [0.902, 0.968] for network A; *r* = 0.84, *p* = 6.5 × 10^−6^, 95% *CI* [0.850, 0.940] for network B), shorter decision times for error trials (*r* = −0.51, *p* = 0.03, 95% *CI* [0.904, 0.967] for network A; *r* = −0.84, *p* = 7.30 × 10^−6^, 95% *CI* [0.953, 0.968] for network B), and decreased decision accuracy (*r* = −0.84, *p* = 6.6 × 10^−6^, 95% *CI* [0.704, 0.783] for network A; *r* = −0.91, *p* = 4.4 × 10^−8^, 95% *CI* [0.784, 0.864] for network B), while decision confidence decreased for correct trials in both networks (*r* = −0.77, *p* = 1.2 × 10^−4^, 95% *CI* [0.950, 0.965] for network A, *r* = −0.73, *p* = 4.3 × 10^−4^, 95% *CI* [0.963, 0.971] for network B) and did not change significantly for error trials in either network (Fig. 5d). These results suggest that, in contrast to anti-phase amplitude modulation, anti-phase frequency modulation exhibited the opposite trend for both network types (enhanced decision accuracy, prolonged correct decision times, shortened error decision times).

**Fig. 5.**
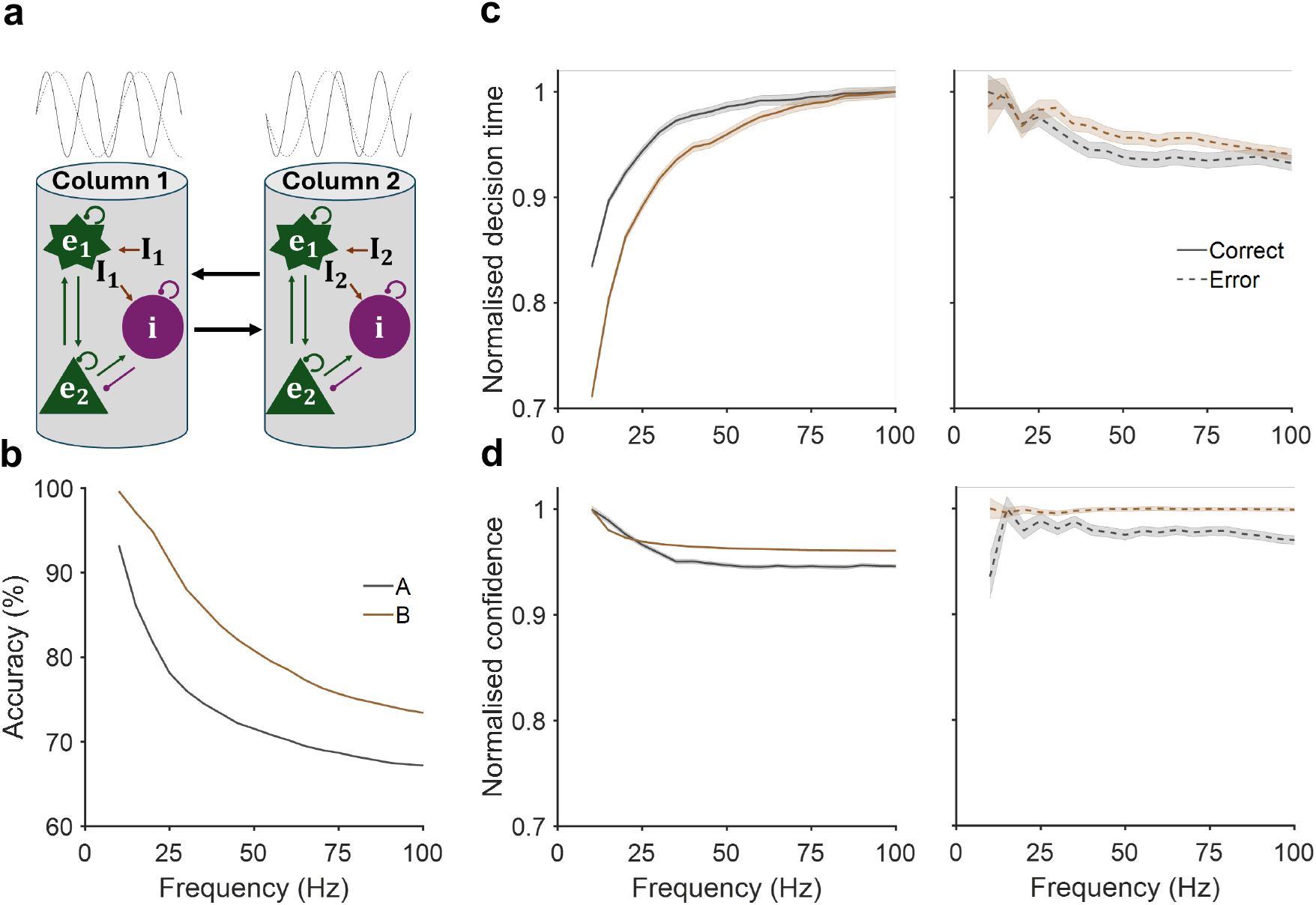
The effects of anti-phase neural oscillatory frequency (amplitude = 0.08 *mA*) are the opposite of those of amplitude oscillation. **a** Anti-phase sinusoidal modulation with varying frequencies applied to both cortical columns for each network. **b** Decision accuracy declined with higher oscillation frequencies in both networks. **c** Normalised decision time increased with increasing amplitude for correct trials (left), while it decreased for error trials (right). **d** Normalised decision confidence decreased for correct trials in both networks, and showed no substantial change for error trials.

### Co-modulation of oscillatory frequency and amplitude on decision-making

We systematically examined the joint influence of oscillatory input frequency and amplitude on decision-making behavior in both Network A and Network B. Decision accuracy was consistently lower across the full amplitude and frequency range for in-phase oscillatory input compared to anti-phase input (Fig. 6).

**Fig. 6.**
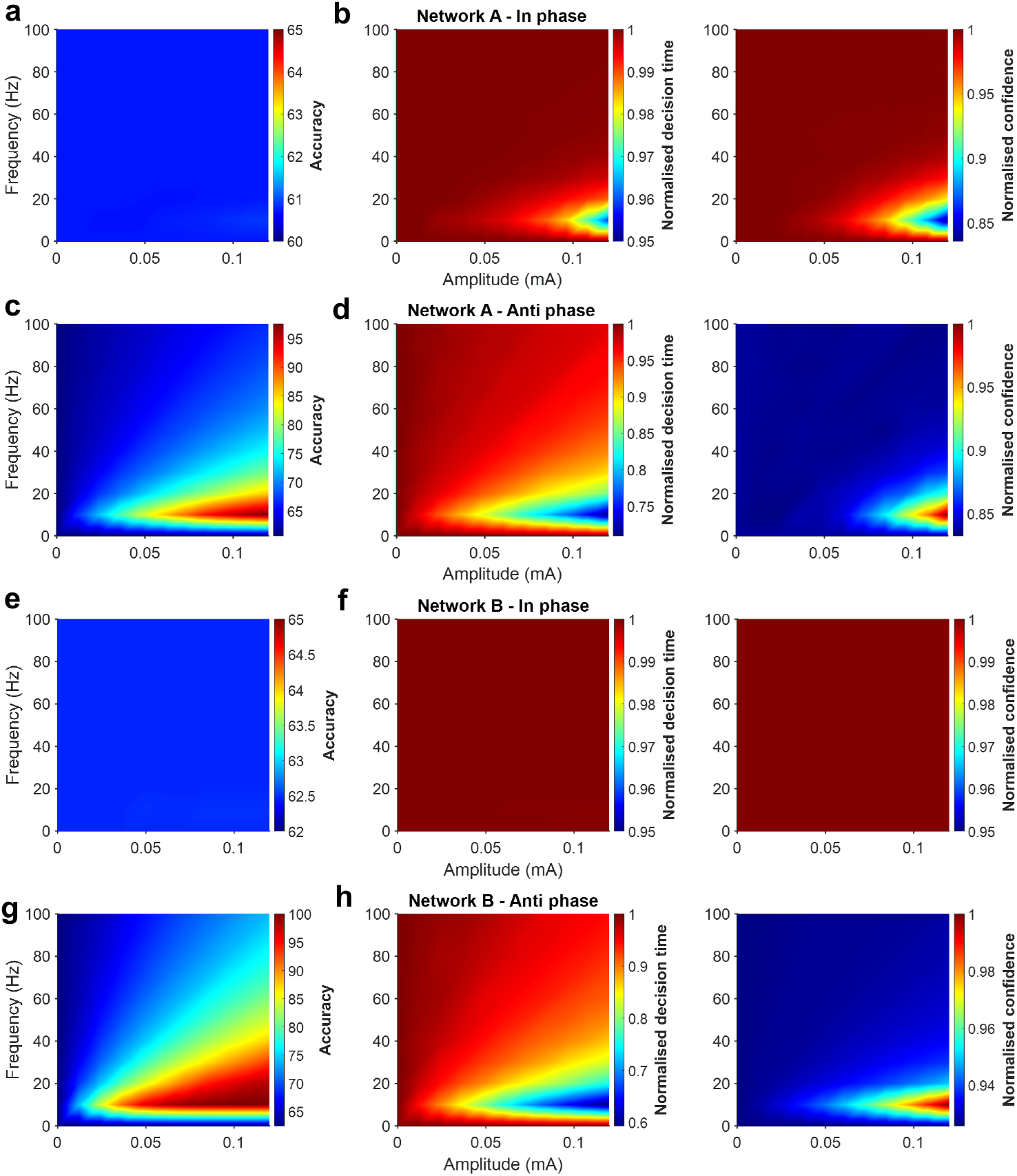
Effects of oscillatory input frequency and amplitude on decision-making behaviour. **Network A (a–d):** Both in-phase and anti-phase oscillations in the theta–beta band (∼4–30 Hz) significantly modulate decision parameters. Under in-phase driving, accuracy remains largely unaffected (a), whereas decision time (b, left) is shortened and confidence (b, right) is diminished across the theta–beta range. Under anti-phase oscillatory input, accuracy improves (c) from theta up to low gamma (∼40 Hz), while decision time decreases (d, left) and confidence increases (d, right) within the theta–beta band. **Network B (e–h):** In-phase oscillations produce minimal changes in accuracy (e), decision time (f, left) or confidence (f, right). By contrast, anti-phase oscillatory input enhances accuracy (g) across the theta to high gamma (∼80 Hz) range and, within the theta–beta band, yields faster decisions (h, left) and higher confidence (h, right).

In Network A, in-phase oscillations left accuracy largely unchanged (Fig. 6a), but were associated with reduced decision time (Fig. 6b, left) and decreased confidence (Fig. 6b, right) across this range. Anti-phase oscillations enhanced accuracy (Fig. 6c) from the theta range up to low gamma (∼40 *Hz*), with the most pronounced improvements occurring in the theta–beta band, where decision time was also reduced (Fig. 6d, left) and confidence increased (6d, right).

In Network B, in-phase oscillatory input exerted little effect on accuracy (Fig. 6e), decision time (Fig. 6f, left), or confidence (Fig. 6f, right) across the examined frequency–amplitude space. By contrast, anti-phase input increased accuracy (Fig. 6g) from theta (∼4 *Hz*) through high gamma (∼80 *Hz*). Within the theta–beta band, these effects were accompanied by shorter decision times (Fig. 6h, left) and higher decision confidence levels (Fig. 6h, right).

In Tables 2–9, partial correlations (Pearson) and p-values are reported for the relationships between oscillatory amplitudes, frequencies, and decision parameters in correct trials, across different ranges of frequencies and amplitudes, respectively. These analyses correspond to Figure 6 and demonstrate the significance of our results across a spectrum of amplitudes and frequencies, rather than for a single specific condition, for both networks under in-phase and anti-phase oscillatory input.

**Table 2:** Partial correlations and p-values for the relationship between the amplitude of in-phase oscillatory input and decision parameters for correct trials across different frequency ranges in Network A. Accuracy (ACC), Decision Time (DT) and Decision Confidence (DC).

| Frequency range (Hz) | Partial correlation | P-value |
| --- | --- | --- |
| 10-20 | ACC: *NaN*<br>DT: $-0.96$<br>DC: $-0.93$ | ACC: *NaN*<br>DT: $8.6 \times 10^{-6}$<br>DC: $1.2 \times 10^{-4}$ |
| 20-30 | ACC: *NaN*<br>DT: $-0.94$<br>DC: $-0.92$ | ACC: *NaN*<br>DT: $6.9 \times 10^{-5}$<br>DC: $1.9 \times 10^{-4}$ |
| 30-40 | ACC: *NaN*<br>DT: $-0.93$<br>DC: $-0.89$ | ACC: *NaN*<br>DT: $8.5 \times 10^{-5}$<br>DC: $6.0 \times 10^{-4}$ |
| 40-50 | ACC: *NaN*<br>DT: $-0.94$<br>DC: $-0.91$ | ACC: *NaN*<br>DT: $5.6 \times 10^{-5}$<br>DC: $3.0 \times 10^{-4}$ |
| 50-60 | ACC: *NaN*<br>DT: $-0.95$<br>DC: $-0.91$ | ACC: *NaN*<br>DT: $2.5 \times 10^{-5}$<br>DC: $2.0 \times 10^{-4}$ |
| 60-70 | ACC: *NaN*<br>DT: $-0.97$<br>DC: $-0.26$ | ACC: *NaN*<br>DT: $4.1 \times 10^{-6}$<br>DC: $4.7 \times 10^{-1}$ |
| 70-80 | ACC: *NaN*<br>DT: $-0.98$<br>DC: $-0.89$ | ACC: *NaN*<br>DT: $6.8 \times 10^{-7}$<br>DC: $5.5 \times 10^{-4}$ |
| 80-90 | ACC: *NaN*<br>DT: $-0.93$<br>DC: $-0.21$ | ACC: *NaN*<br>DT: $8.4 \times 10^{-5}$<br>DC: $5.6 \times 10^{-1}$ |
| 90-100 | ACC: *NaN*<br>DT: $-0.99$<br>DC: $-0.90$ | ACC: *NaN*<br>DT: $2.4 \times 10^{-8}$<br>DC: $4.0 \times 10^{-4}$ |

**Table 3:** Partial correlations and p-values for the relationship between the frequency of in-phase oscillatory input and decision parameters for correct trials across different amplitude ranges in Network A. Accuracy (ACC), Decision Time (DT) and Decision Confidence (DC).

| Amplitude range (mA) | Partial correlation | P-value |
| --- | --- | --- |
| 0.012-0.024 | ACC: *NaN*<br>DT: 0.56<br>DC: 0.42 | ACC: *NaN*<br>DT: $9.1 \times 10^{-2}$<br>DC: $2.3 \times 10^{-1}$ |
| 0.024-0.036 | ACC: *NaN*<br>DT: 0.63<br>DC: 0.61 | ACC: *NaN*<br>DT: $5.3 \times 10^{-2}$<br>DC: $5.9 \times 10^{-2}$ |
| 0.036-0.048 | ACC: *NaN*<br>DT: 0.70<br>DC: 0.68 | ACC: *NaN*<br>DT: $2.0 \times 10^{-2}$<br>DC: $3.1 \times 10^{-2}$ |
| 0.048-0.060 | ACC: *NaN*<br>DT: 0.73 | ACC: *NaN*<br>DT: $2.0 \times 10^{-2}$ |
| | DC: 0.70 | DC: $2.5 \times 10^{-2}$ |
| 0.060-0.072 | ACC: *NaN*<br>DT: 0.73<br>DC: 0.72 | ACC: *NaN*<br>DT: $1.5 \times 10^{-2}$<br>DC: $1.8 \times 10^{-2}$ |
| 0.072-0.084 | ACC: *NaN*<br>DT: 0.73<br>DC: 0.72 | ACC: *NaN*<br>DT: $1.7 \times 10^{-2}$<br>DC: $1.8 \times 10^{-2}$ |
| 0.084-0.096 | ACC: *NaN*<br>DT: 0.72<br>DC: 0.72 | ACC: *NaN*<br>DT: $1.9 \times 10^{-2}$<br>DC: $1.8 \times 10^{-2}$ |
| 0.096-0.108 | ACC: *NaN*<br>DT: 0.69<br>DC: 0.71 | ACC: *NaN*<br>DT: $2.6 \times 10^{-2}$<br>DC: $2.0 \times 10^{-2}$ |
| 0.108-0.120 | ACC: *NaN*<br>DT: 0.67<br>DC: 0.71 | ACC: *NaN*<br>DT: $3.2 \times 10^{-2}$<br>DC: $2.0 \times 10^{-2}$ |

**Table 4:** Partial correlations and p-values for the relationship between the amplitude of anti-phase oscillatory input and decision parameters for correct trials across different frequency ranges in Network A. Accuracy (ACC), Decision Time (DT) and Decision Confidence (DC).

| Frequency range (Hz) | Partial correlation | P-value |
| --- | --- | --- |
| 10-20 | ACC: 0.99<br>DT: -0.99<br>DC: 0.94 | ACC: $1.6 \times 10^{-10}$<br>DT: $1.3 \times 10^{-13}$<br>DC: $6.3 \times 10^{-5}$ |
| | ACC: 0.99 | ACC: $2.1 \times 10^{-12}$ |
| 20-30 | DT: $-0.99$<br>DC: 0.92 | DT: $2.0 \times 10^{-12}$<br>DC: $1.4 \times 10^{-4}$ |
| 30-40 | ACC: 0.99<br>DT: $-0.99$<br>DC: 0.89 | ACC: $7.0 \times 10^{-12}$<br>DT: $9.2 \times 10^{-12}$<br>DC: $6.0 \times 10^{-4}$ |
| 40-50 | ACC: 0.99<br>DT: $-0.99$<br>DC: 0.73 | ACC: $3.0 \times 10^{-12}$<br>DT: $5.4 \times 10^{-11}$<br>DC: $2.0 \times 10^{-2}$ |
| 50-60 | ACC: 0.99<br>DT: $-0.99$<br>DC: 0.78 | ACC: $3.1 \times 10^{-12}$<br>DT: $9.2 \times 10^{-11}$<br>DC: $7.0 \times 10^{-3}$ |
| 60-70 | ACC: 0.99<br>DT: $-0.99$<br>DC: 0.11 | ACC: $9.1 \times 10^{-13}$<br>DT: $1.6 \times 10^{-8}$<br>DC: $7.6 \times 10^{-1}$ |
| 70-80 | ACC: 0.99<br>DT: $-0.99$<br>DC: $-0.02$ | ACC: $1.2 \times 10^{-13}$<br>DT: $1.1 \times 10^{-8}$<br>DC: $9.6 \times 10^{-1}$ |
| 80-90 | ACC: 0.99<br>DT: $-0.99$<br>DC: $-0.49$ | ACC: $7.3 \times 10^{-13}$<br>DT: $1.5 \times 10^{-8}$<br>DC: $1.5 \times 10^{-2}$ |
| 90-100 | ACC: 0.99<br>DT: $-0.99$<br>DC: $-0.48$ | ACC: $2.0 \times 10^{-12}$<br>DT: $1.8 \times 10^{-8}$<br>DC: $1.6 \times 10^{-2}$ |

**Table 5:** Partial correlations and p-values for the relationship between the frequency of anti-phase oscillatory input and decision parameters for correct trials across different amplitude ranges in Network A. Accuracy (ACC), Decision Time (DT) and Decision Confidence (DC).

| Amplitude range (mA) | Partial correlation | P-value |
| --- | --- | --- |
| 0.012-0.024 | ACC: $-0.82$<br>DT: $0.81$<br>DC: $0.61$ | ACC: $3.0 \times 10^{-3}$<br>DT: $5.0 \times 10^{-3}$<br>DC: $6.0 \times 10^{-2}$ |
| 0.024-0.036 | ACC: $-0.83$<br>DT: $0.80$<br>DC: $-0.46$ | ACC: $3.0 \times 10^{-3}$<br>DT: $6.0 \times 10^{-3}$<br>DC: $1.8 \times 10^{-2}$ |
| 0.036-0.048 | ACC: $-0.84$<br>DT: $0.81$<br>DC: $-0.81$ | ACC: $2.0 \times 10^{-3}$<br>DT: $5.0 \times 10^{-3}$<br>DC: $5.0 \times 10^{-3}$ |
| 0.048-0.060 | ACC: $-0.84$<br>DT: $0.82$<br>DC: $-0.75$ | ACC: $2.0 \times 10^{-3}$<br>DT: $4.0 \times 10^{-3}$<br>DC: $1.0 \times 10^{-2}$ |
| 0.060-0.072 | ACC: $-0.85$<br>DT: $0.82$<br>DC: $-0.72$ | ACC: $2.0 \times 10^{-3}$<br>DT: $3.0 \times 10^{-3}$<br>DC: $2.0 \times 10^{-2}$ |
| 0.072-0.084 | ACC: $-0.86$<br>DT: $0.83$<br>DC: $-0.74$ | ACC: $1.0 \times 10^{-3}$<br>DT: $3.0 \times 10^{-3}$<br>DC: $1.0 \times 10^{-2}$ |
| 0.084-0.096 | ACC: $-0.87$<br>DT: $-0.99$<br>DC: $0.83$ | ACC: $1.0 \times 10^{-3}$<br>DT: $1.1 \times 10^{-8}$ |
| | | DC: $3.0 \times 10^{-3}$ |
| 0.096-0.108 | ACC: $-0.88$<br>DT: $0.82$<br>DC: $-0.77$ | ACC: $8.2 \times 10^{-4}$<br>DT: $3.0 \times 10^{-3}$<br>DC: $9.0 \times 10^{-3}$ |
| 0.108-0.120 | ACC: $-0.89$<br>DT: $0.82$<br>DC: $-0.75$ | ACC: $5.0 \times 10^{-4}$<br>DT: $3.0 \times 10^{-3}$<br>DC: $1.0 \times 10^{-2}$ |

**Table 6:** Partial correlations and p-values for the relationship between the amplitude of in-phase oscillatory input and decision parameters for correct trials across different frequency ranges in Network B. Accuracy (ACC), Decision Time (DT) and Decision Confidence (DC).

| Frequency range (Hz) | Partial correlation | P-value |
| --- | --- | --- |
| 10-20 | ACC: *NaN*<br>DT: *NaN*<br>DC: *NaN* | ACC: *NaN*<br>DT: *NaN*<br>DC: *NaN* |
| 20-30 | ACC: *NaN*<br>DT: *NaN*<br>DC: *NaN* | ACC: *NaN*<br>DT: *NaN*<br>DC: *NaN* |
| 30-40 | ACC: *NaN*<br>DT: *NaN*<br>DC: *NaN* | ACC: *NaN*<br>DT: *NaN*<br>DC: *NaN* |
| 40-50 | ACC: *NaN*<br>DT: *NaN*<br>DC: *NaN* | ACC: *NaN*<br>DT: *NaN*<br>DC: *NaN* |
| 50-60 | ACC: *NaN*<br>DT: *NaN*<br>DC: *NaN* | ACC: *NaN*<br>DT: *NaN*<br>DC: *NaN* |
| 60-70 | ACC: *NaN*<br>DT: *NaN*<br>DC: *NaN* | ACC: *NaN*<br>DT: *NaN*<br>DC: *NaN* |
| 70-80 | ACC: *NaN*<br>DT: *NaN*<br>DC: *NaN* | ACC: *NaN*<br>DT: *NaN*<br>DC: *NaN* |
| 80-90 | ACC: *NaN*<br>DT: *NaN*<br>DC: *NaN* | ACC: *NaN*<br>DT: *NaN*<br>DC: *NaN* |
| 90-100 | ACC: *NaN*<br>DT: *NaN*<br>DC: *NaN* | ACC: *NaN*<br>DT: *NaN*<br>DC: *NaN* |

**Table 7:** Partial correlations and p-values for the relationship between the frequency of in-phase oscillatory input and decision parameters for correct trials across different amplitude ranges in Network B. Accuracy (ACC), Decision Time (DT) and Decision Confidence (DC).

| Amplitude range (mA) | Partial correlation | P-value |
| --- | --- | --- |
| 0.012-0.024 | ACC: *NaN*<br>DT: *NaN*<br>DC: *NaN* | ACC: *NaN*<br>DT: *NaN*<br>DC: *NaN* |
| 0.024-0.036 | ACC: *NaN*<br>DT: *NaN* | ACC: *NaN*<br>DT: *NaN* |
|  | DC: *NaN* | DC: *NaN* |
| 0.036-0.048 | ACC: *NaN*<br>DT: *NaN*<br>DC: *NaN* | ACC: *NaN*<br>DT: *NaN*<br>DC: *NaN* |
| 0.048-0.060 | ACC: *NaN*<br>DT: *NaN*<br>DC: *NaN* | ACC: *NaN*<br>DT: *NaN*<br>DC: *NaN* |
| 0.060-0.072 | ACC: *NaN*<br>DT: *NaN*<br>DC: *NaN* | ACC: *NaN*<br>DT: *NaN*<br>DC: *NaN* |
| 0.072-0.084 | ACC: *NaN*<br>DT: *NaN*<br>DC: *NaN* | ACC: *NaN*<br>DT: *NaN*<br>DC: *NaN* |
| 0.084-0.096 | ACC: *NaN*<br>DT: *NaN*<br>DC: *NaN* | ACC: *NaN*<br>DT: *NaN*<br>DC: *NaN* |
| 0.096-0.108 | ACC: *NaN*<br>DT: *NaN*<br>DC: *NaN* | ACC: *NaN*<br>DT: *NaN*<br>DC: *NaN* |
| 0.108-0.120 | ACC: *NaN*<br>DT: *NaN*<br>DC: *NaN* | ACC: *NaN*<br>DT: *NaN*<br>DC: *NaN* |

**Table 8:**
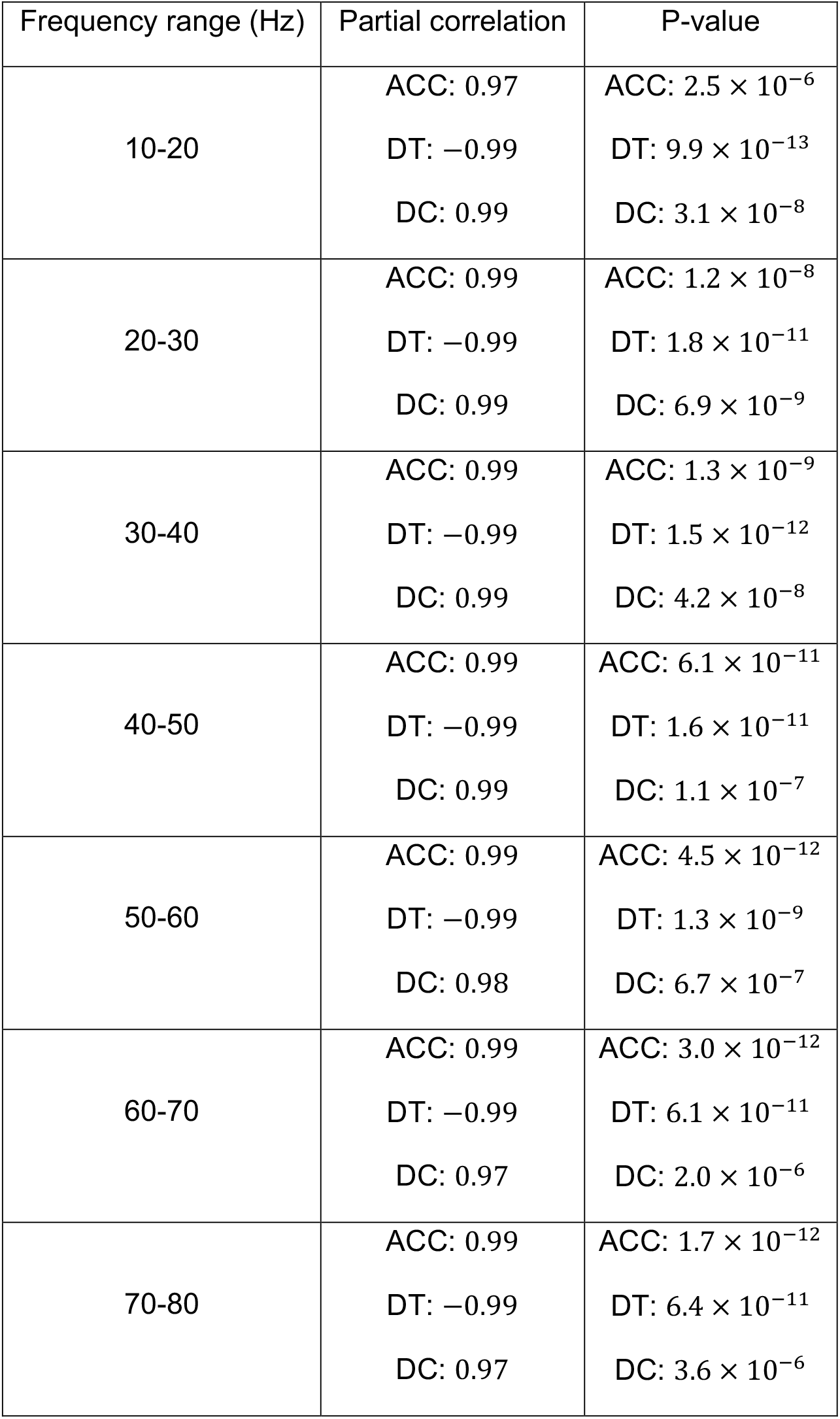
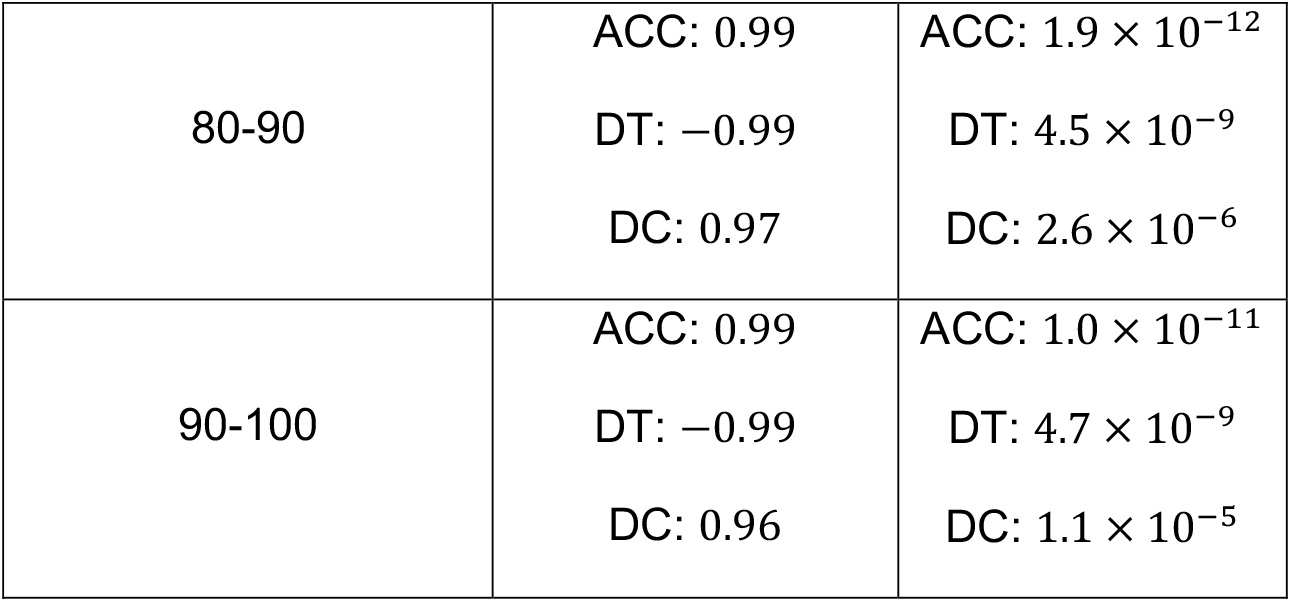
Partial correlations and p-values for the relationship between the amplitude of anti-phase oscillatory input and decision parameters for correct trials across different frequency ranges in Network B. Accuracy (ACC), Decision Time (DT) and Decision Confidence (DC).

| Frequency range (Hz) | Partial correlation | P-value |
| --- | --- | --- |
| 10-20 | ACC: 0.97<br>DT: -0.99<br>DC: 0.99 | ACC: $2.5 \times 10^{-6}$<br>DT: $9.9 \times 10^{-13}$<br>DC: $3.1 \times 10^{-8}$ |
| 20-30 | ACC: 0.99<br>DT: -0.99<br>DC: 0.99 | ACC: $1.2 \times 10^{-8}$<br>DT: $1.8 \times 10^{-11}$<br>DC: $6.9 \times 10^{-9}$ |
| 30-40 | ACC: 0.99<br>DT: -0.99<br>DC: 0.99 | ACC: $1.3 \times 10^{-9}$<br>DT: $1.5 \times 10^{-12}$<br>DC: $4.2 \times 10^{-8}$ |
| 40-50 | ACC: 0.99<br>DT: -0.99<br>DC: 0.99 | ACC: $6.1 \times 10^{-11}$<br>DT: $1.6 \times 10^{-11}$<br>DC: $1.1 \times 10^{-7}$ |
| 50-60 | ACC: 0.99<br>DT: -0.99<br>DC: 0.98 | ACC: $4.5 \times 10^{-12}$<br>DT: $1.3 \times 10^{-9}$<br>DC: $6.7 \times 10^{-7}$ |
| 60-70 | ACC: 0.99<br>DT: -0.99<br>DC: 0.97 | ACC: $3.0 \times 10^{-12}$<br>DT: $6.1 \times 10^{-11}$<br>DC: $2.0 \times 10^{-6}$ |
| 70-80 | ACC: 0.99<br>DT: -0.99<br>DC: 0.97 | ACC: $1.7 \times 10^{-12}$<br>DT: $6.4 \times 10^{-11}$<br>DC: $3.6 \times 10^{-6}$ |
| 80-90 | ACC: 0.99<br>DT: -0.99<br>DC: 0.97 | ACC: $1.9 \times 10^{-12}$<br>DT: $4.5 \times 10^{-9}$<br>DC: $2.6 \times 10^{-6}$ |
| 90-100 | ACC: 0.99<br>DT: -0.99<br>DC: 0.96 | ACC: $1.0 \times 10^{-11}$<br>DT: $4.7 \times 10^{-9}$<br>DC: $1.1 \times 10^{-5}$ |

**Table 9:** Partial correlations and p-values for the relationship between the frequency of anti-phase oscillatory input and decision parameters for correct trials across different amplitude ranges in Network B. Accuracy (ACC), Decision Time (DT) and Decision Confidence (DC).

| Amplitude range (mA) | Partial correlation | P-value |
| --- | --- | --- |
| 0.012-0.024 | ACC: -0.84<br>DT: 0.81<br>DC: -0.73 | ACC: $2.0 \times 10^{-3}$<br>DT: $4.2 \times 10^{-3}$<br>DC: $1.6 \times 10^{-2}$ |
| 0.024-0.036 | ACC: -0.86<br>DT: 0.84<br>DC: -0.72 | ACC: $1.5 \times 10^{-3}$<br>DT: $2.5 \times 10^{-3}$<br>DC: $2.0 \times 10^{-2}$ |
| 0.036-0.048 | ACC: -0.88<br>DT: 0.84<br>DC: -0.73 | ACC: $8.0 \times 10^{-4}$<br>DT: $2.4 \times 10^{-3}$<br>DC: $1.7 \times 10^{-2}$ |
| 0.048-0.060 | ACC: -0.90<br>DT: 0.83<br>DC: -0.74 | ACC: $3.2 \times 10^{-4}$<br>DT: $3.0 \times 10^{-3}$<br>DC: $1.5 \times 10^{-2}$ |
| | ACC: -0.93 | ACC: $1.0 \times 10^{-4}$ |
| 0.060-0.072 | DT: 0.82<br>DC: -0.74 | DT: $3.3 \times 10^{-3}$<br>DC: $1.5 \times 10^{-2}$ |
| 0.072-0.084 | ACC: -0.95<br>DT: 0.83<br>DC: -0.73 | ACC: $3.1 \times 10^{-5}$<br>DT: $3.1 \times 10^{-3}$<br>DC: $1.7 \times 10^{-2}$ |
| 0.084-0.096 | ACC: -0.96<br>DT: 0.84<br>DC: -0.72 | ACC: $8.7 \times 10^{-6}$<br>DT: $2.3 \times 10^{-3}$<br>DC: $1.8 \times 10^{-2}$ |
| 0.096-0.108 | ACC: -0.97<br>DT: 0.85<br>DC: -0.72 | ACC: $3.1 \times 10^{-6}$<br>DT: $1.9 \times 10^{-3}$<br>DC: $2.0 \times 10^{-2}$ |
| 0.108-0.120 | ACC: -0.98<br>DT: 0.86<br>DC: -0.71 | ACC: $8.0 \times 10^{-7}$<br>DT: $1.5 \times 10^{-3}$<br>DC: $2.0 \times 10^{-2}$ |

These findings align with prior experiments, showing that alpha rhythms (Samaha et al., 2017, 2020) and beta–gamma band activity (Donner et al., 2009) play key roles in perceptual decision-making.

### Oscillatory phase difference progressively affects decision behaviours

Given the prominent effects by alpha band frequency, we next investigated how gradual changes in its oscillatory phase differences (Δ*ϕ*) between neural inputs affect decision-making in networks A and B. Building on prior analyses of in-phase and anti-phase effects, we adjusted Δ*ϕ* between inputs to network A, with the phase to column 1 (encoding correct decision) leading that of column 2 (encoding error decision) (Fig. S3). In network A, decision accuracy increased sharply at larger phase differences (e.g., Δ*ϕ* = π) compared to smaller ones (e.g., Δ*ϕ* = 0), especially as oscillatory amplitude increased (Fig. S3a). Conversely, decision time decreased more rapidly at larger Δ*ϕ* (Fig. S3b).

Interestingly, as phase difference increased, decision confidence changed from decreasing to increasing trends with oscillation amplitude (Fig. S3c). Network B exhibited similar trends for decision accuracy and decision time, but decision confidence only (slightly) increased with oscillation amplitude (Fig. S3d–f).

When the oscillatory input to column 2 (error decision) led column 1 (correct decision), network A exhibited a decline in decision accuracy as amplitude increased at larger Δ*ϕ* (Fig. S4a). As the phase difference increased, decision time and decision confidence (**c**) changed from decreasing to increasing trends with oscillation amplitude (Fig. S4b−c). For network B, increasing Δ*ϕ* led to decreased decision accuracy as in network A, but decision time only increased with oscillation amplitude while decision confidence stayed largely constant (Fig. S4d−f).

After reversing the phase lead between columns 1 and 2, for network A, as Δ*ϕ* increased, decision accuracy progressively increased but with decreasing trend with oscillation frequency (Fig. S5a), while decision time decreased but with increasing trend with oscillation frequency (Fig. S5b). The decision confidence changed from increasing to decreasing trends with oscillation frequency (Fig. S5c). For network B, decision accuracy and decision time exhibited similar trends as in network A, but decision confidence only slightly decreased with oscillation frequency (Fig. S5d−f).

With the input to column 2 leading again, increasing Δ*ϕ* led to decision accuracy progressively decreasing with increasing trend with oscillation frequency (Fig. S6a), and decision time decreased with oscillation frequency (Fig. S6b), while decision confidence changed from increasing to decreasing trends with oscillation frequency (Fig. 6c). For network B, the trends are similar except that decision confidence did not significantly change with oscillation frequency or phase difference (Fig. S6d-f).

Overall, we have shown that where the oscillatory phase lead resided, in neural population encoding correct or error decision, can have decision network be differently modulated by phase difference and oscillation amplitude or frequency. The non-monotonic effects on decision confidence deserve further discussion. Specifically, as the model’s decision confidence is defined as the difference between the winning and losing population’s EPSP activities at the moment of decision threshold crossing, and given that network A is more sensitive to oscillatory influences than network B, when the phase difference between the oscillatory inputs to both columns is closer to π, the difference between the winner and loser EPSP activities reached its maximum value, whereas at a phase difference closer to 0, this difference reached its minimum value. Moreover, depending on the interplay between stimulus input, noise and intrinsic network dynamics, the decision confidence trends of network A could reverse. Thus, our simulation results highlight the intricate role of phase relationships in neural oscillatory modulations on decision-making.

To further investigate how the timing of sensory input interacts with inter-columnar oscillatory phase, we systematically varied both the relative phase of stimulus onset (*φ*_stim_) and the phase difference between the two cortical columns (Δ*φ*). The stimulation phase was referenced to column 1. For each combination, we measured decision accuracy, normalised decision time, and normalised decision confidence.

When *φ*_stim_ was fixed at 0, increasing (Δ*φ*) from 0 to π yielded a monotonic improvement in accuracy (Fig. 7a−b), a corresponding decrease in decision time (Fig. 7c-d), and an increase in confidence (Fig. 7d-f). Extending this analysis across other *φ*_stim_ values revealed a clear phase-dependent modulation: certain combinations of *φ*_stim_ and Δ*φ* selectively enhanced performance metrics, while others diminished them (Fig. 7). These results demonstrate that decision outputs are shaped not only by inter-columnar phase relationships but also by the precise timing of sensory inputs relative to ongoing oscillations.

**Fig. 7.**
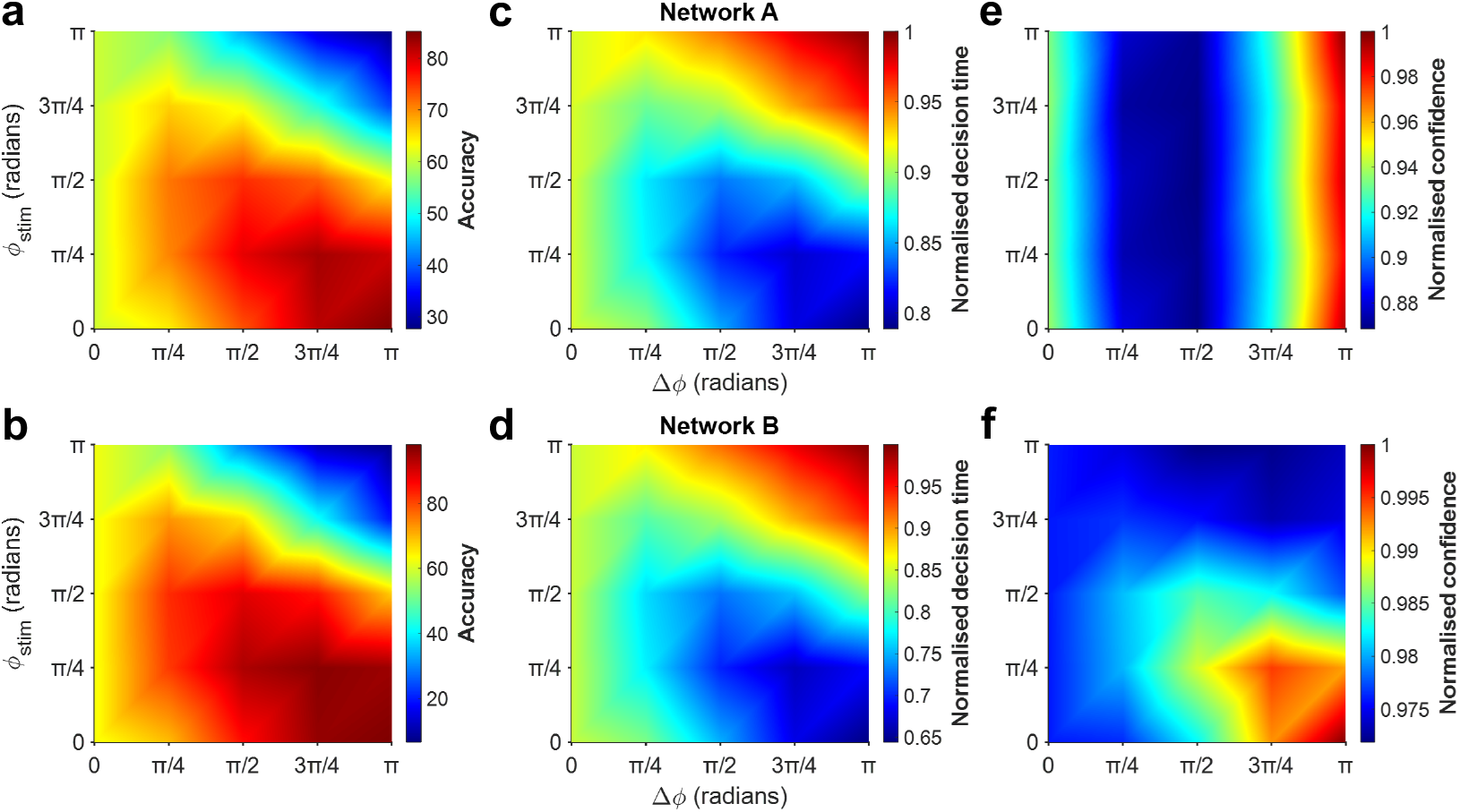
Phase-dependent effects of sensory input within alpha band frequency. Each panel plots the relative stimulus phase (*φ*_stim_) against the inter-column phase difference (Δ*φ*) for decision accuracy (**a, b**), normalised decision time (**c, d**) and normalised decision confidence (**e, f**). Consistent with our earlier results, when the stimulus phase is fixed at 0 and the inter-column phase difference is increased from 0 to π, we observe: (**a, b**) higher decision accuracy, (**c, d**) shorter decision times, and (**e, f**) greater decision confidence.

These phase-dependent maps (Fig. 7) show a clear dip in decision confidence in Network A when the phase difference is about one-quarter to one-half of a cycle. This suggests that small intercolumnar delays of ∼20–40 ms can noticeably lower confidence, regardless of the stimulus phase.

### In-phase modulation affects excitatory neural populations, while anti-phase affects inhibitory neural populations

So far, we have applied oscillatory inputs to both excitatory and inhibitory neural populations. Hence, to understand the contributions of each neural population, we examined the behavioural effects of selectively applying oscillatory inputs to excitatory or inhibitory neural populations in networks with distinct intrinsic timescales. Given the prominent effects of ∼12 *Hz* modulatory effects (Fig. 6), we shall henceforth focus on this frequency when varying oscillation amplitude.

When oscillatory input was applied exclusively to only excitatory neural populations, in-phase stimulation at 12 *Hz* (Fig. 8a) produced no change in decision accuracy (Fig. 8b) across amplitude variations in either network. In Network A, decision time decreased for correct trials (Fig. 8c, left; *r* = −0.91, *p* = 6.2 × 10^−9^, 95% *CI* [0.989, 0.995]) and remained nearly unchanged for error trials (Fig. 8c, right), accompanied by reduced decision confidence for correct (Fig. 8d, left; *r* = −0.90, *p* = 2.7 × 10^−8^, 95% *CI* [0.948, 0.982]) and error trials (Fig. 8d, right; *r* = −0.78, *p* = 2.5 × 10^−5^, 95% *CI* [0.990, 0.997]), whereas Network B showed no significant changes in either measure (Fig. 8b-d).

**Fig. 8.**
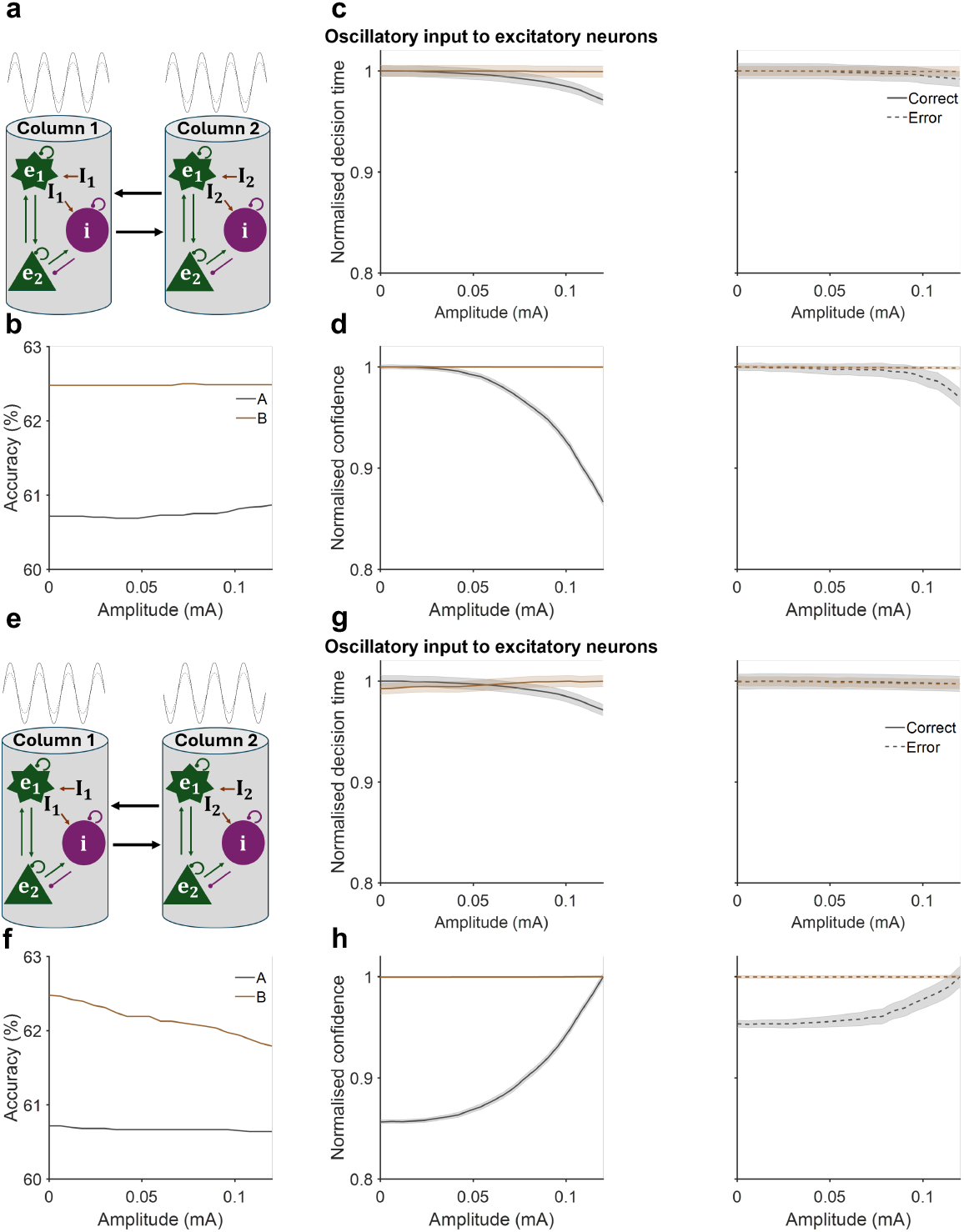
Effects of in-phase and anti-phase 12 Hz neural oscillation amplitude of input to excitatory neurons in networks with differing timescales. **a** Illustration of in-phase sinusoidal modulation at varying amplitudes applied simultaneously to both cortical columns and exclusively to excitatory neurons in each network. **b** Decision accuracy remained stable across amplitude variations for both networks. **c** Network A (grey) exhibited reduced normalised decision time with increasing oscillation amplitude for correct trials (left), while decision time remained nearly unchanged for error trials (right). Network B (brown) showed no significant change in normalised decision time for either trial type. **d** Network A demonstrated decreased normalised decision confidence with increasing oscillation amplitude. In contrast, Network B displayed no significant change in decision confidence for either correct (left) or error (right) trials. **e** Illustration of anti-phase sinusoidal modulation at varying amplitudes applied simultaneously to both cortical columns and exclusively to excitatory neurons in each network. **f** Decision accuracy decreased with higher oscillation amplitudes in Network B and remained constant in Network A. **g** Normalised decision time decreased with increasing oscillation amplitude for correct trials of Network A and remained constant for Network B (left). For error trials, normalised decision time remained constant across both networks (right). **h** Normalised decision confidence increased in Network A for both correct and error trials, whereas it remained constant in Network B for both trial types.

Anti-phase input (Fig. 8e) left accuracy unchanged in Network A but slightly reduced it in Network B (Fig. 8f; *r* = −0.99, *p* = 3.3 × 10^−18^, 95% *CI* [0.620, 0.622]). In Network A, higher amplitudes reduced decision time for correct trials (Fig. 8g, left; *r* = −0.91, *p* = 1.0 × 10^−8^, 95% *CI* [0.989, 0.996]) and unchanged for error trials (Fig. 8g, right) and increased confidence across trial types (Fig. 8h; *r* = 0.92, *p* = 3.6 × 10^−9^, 95% *CI* [0.878, 0.916] for correct trials; *r* = 0.87, *p* = 2.3 × 10^−7^, 95% *CI* [0.958, 0.970] for error trials), while Network B remained largely unaffected in decision time and confidence (Fig. 8f-h).

Frequency modulation of in-phase excitation (Fig. 9a) did not alter accuracy in either network (Fig. 9b). In Network A, it slightly increased decision time for correct trials (Fig. 9c, left; *r* = 0.73, *p* = 3.3 × 10^−4^, 95% *CI* [0.996, 0.999]) while leaving error trials unaffected (Fig. 9c, right), whereas in Network B, decision time remained unchanged for both trial types (Fig. 9c). Decision confidence in Network A increased for correct trials (*r* = 0.71, *p* = 6.9 × 10^−4^, 95% *CI* [0.985, 0.997]) but did not change significantly for error trials, whereas it was unaffected in Network B (Fig. 9d). Under anti-phase excitation (Fig. 9e), increasing frequency in Network A did not change accuracy (Fig. 9f) or decision time for either correct (Fig. 9g, left) or error trials (Fig. 9g, right), but it reduced confidence for both trials (Fig. 9h; *r* = −0.80, *p* = 4.4 × 10^−5^, 95% *CI* [0.953, 0.966] for correct trials; *r* =−0.76, *p* = 1.8 × 10^−4^, 95% *CI* [0.980, 0.985] for error trials). In contrast, Network B showed no substantial changes in any decision parameters (Fig. 9f–h).

**Fig. 9.**
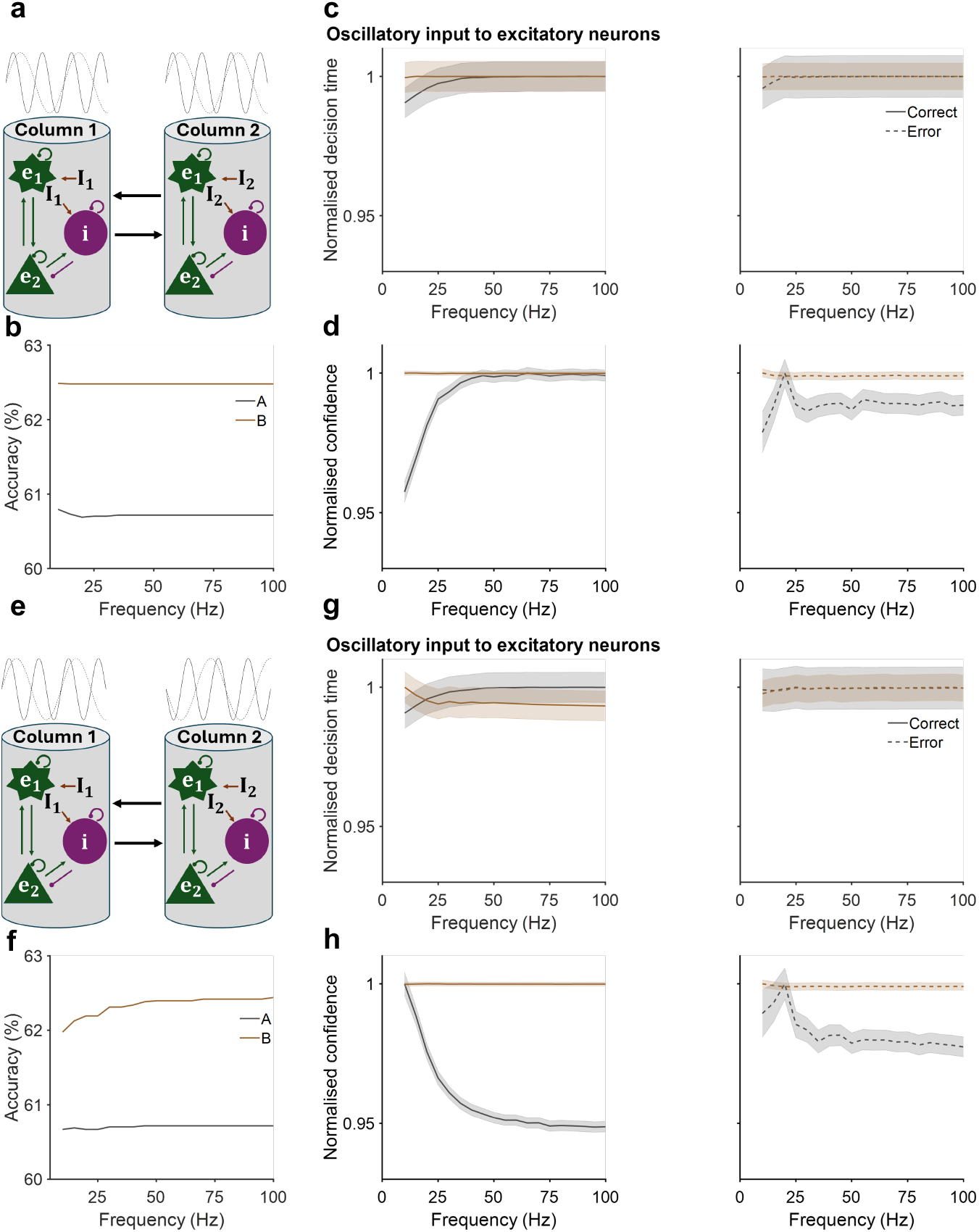
Frequency-dependent effects of in-phase and anti-phase neural oscillatory input to excitatory neurons (amplitude = 0.08 mA) on networks with distinct time constants. **a** Schematic of in-phase sinusoidal modulation applied at varying frequencies to both cortical columns and exclusively to excitatory neurons in each network. **b** Decision accuracy remained unchanged across frequencies for both networks. **c, d** In Network A (grey), normalised decision time (**c**) and decision confidence (**d**) increased for correct (left) and did not change significantly for error trials (right). Network B (brown) showed no significant changes in either metric for both correct and error trials. **e** Schematic of anti-phase sinusoidal modulation at varying frequencies applied to both cortical columns and exclusively to excitatory neurons. **f** Decision accuracy remained nearly constant as oscillation frequency increased in both networks. **g** Normalised decision time remained essentially constant for both correct (left) and error (right) trials in both networks. **h** Normalised decision confidence decreased for correct trials in Network A; for error trials it also declined in Network A but remained largely unchanged in Network B for both trial types.

When oscillatory input was applied exclusively to inhibitory neural populations, in-phase stimulation had minimal effects on decision variables across different amplitudes in both networks (Fig. 10a–d). In contrast, anti-phase input (Fig. 10e) at 12 *Hz* increased accuracy (Fig. 10f; *r* = 0.99, *p* = 8.8 × 10^−18^, 95% *CI* [0.762, 0.852] for Network A; *r* = 0.89, *p* = 4.6 × 10^−8^, 95% *CI* [0.851, 0.950] for Network B) and reduced decision time for correct trials in both networks (Fig. 10g, left; *r* = −0.99, *p* = 1.4 × 10^−25^, 95% *CI* [0.860, 0.913] for Network A; *r* = −0.99, *p* = 1.8 × 10^−22^, 95% *CI* [0.751, 0.845] for Network B), while leaving confidence largely unchanged in Network A and slightly increasing it in Network B (Fig. 10h, left; *r* = 0.98, *p* = 3.2 × 10^−15^, 95% *CI* [0.958, 0.973] for Network B). For error trials, increasing amplitude prolonged decision time (Fig. 10g, right) in both networks (*r* = 0.99, *p* = 2.5 × 10^−16^, 95% *CI* [0.917, 0.955] for Network A; *r* = 0.91, *p* = 1.8 × 10^−8^, 95% *CI* [0.803, 0.859] for Network B) without substantially affecting confidence (Fig. 10h, right).

**Fig. 10.**
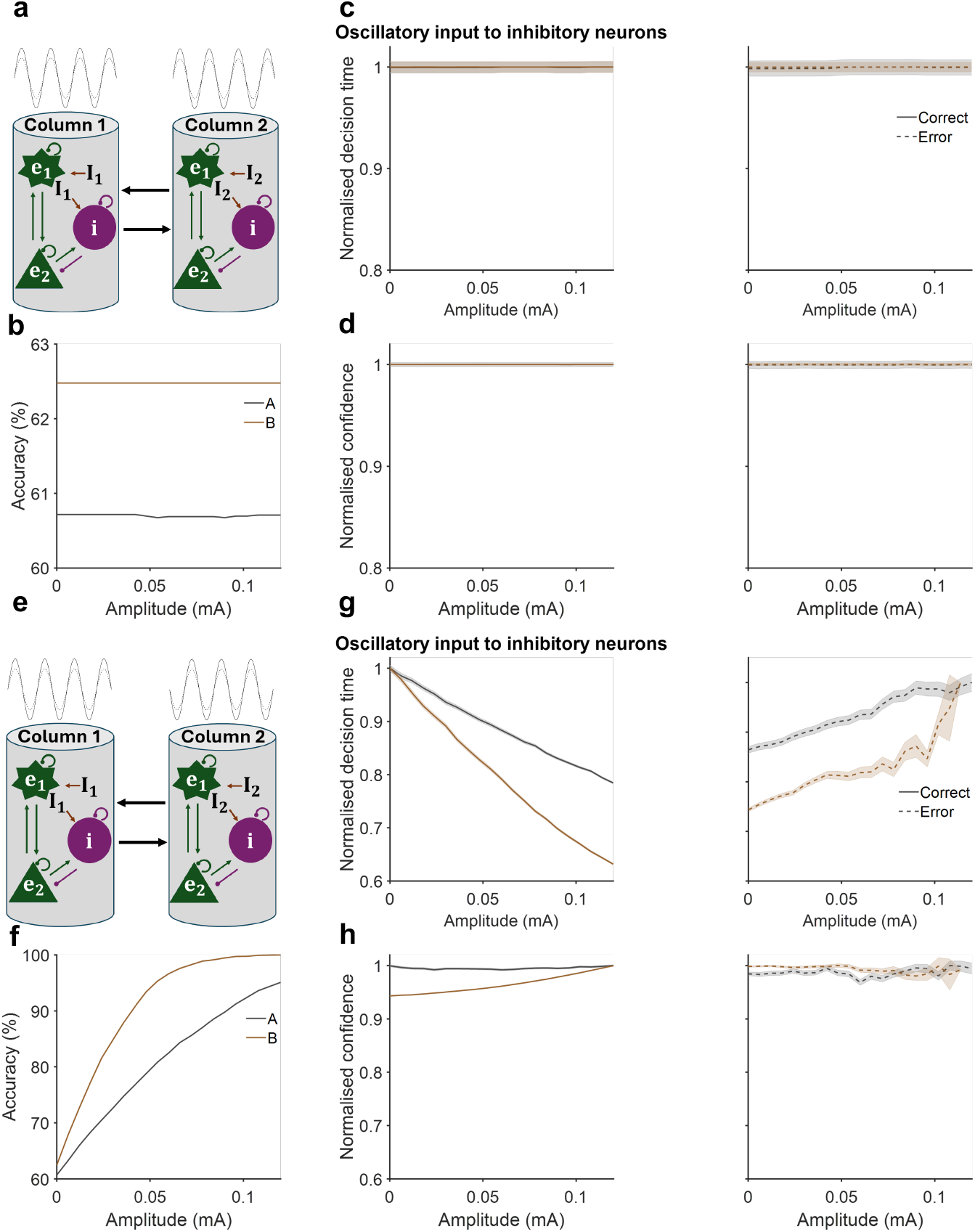
Effects of in-phase and anti-phase 12 Hz neural oscillation amplitude of input to inhibitory neurons in networks with differing timescales. **a** Illustration of in-phase sinusoidal modulation at varying amplitudes applied simultaneously to both cortical columns and exclusively to inhibitory neurons in each network. **b** Decision accuracy remained stable across amplitude variations for both networks. **c, d** Normalised decision time and confidence remained unchanged with increasing oscillation amplitude for both correct (left) and error (right) trials in both networks. **e** Illustration of anti-phase sinusoidal modulation at varying amplitudes applied simultaneously to both cortical columns and exclusively to inhibitory neurons in each network. **f** Decision accuracy increased with higher oscillation amplitudes in both Network A and Network B. **g** Normalised decision time decreased with increasing amplitude for correct trials in both networks (left), whereas for error trials it increased in both networks (right). **h** Normalised decision confidence increased in Network B for correct trials and remained constant for error trials; in Network A, it remained constant for both trial types.

Applying oscillatory input solely to inhibitory neural populations produced minimal changes in decision variables under in-phase stimulation across all tested frequencies in both networks (Fig. 11a–d). Under anti-phase inhibition (Fig. 11e), increasing frequency reduced accuracy (Fig. 11f; *r* = −0.86, *p* = 2.1 × 10^−6^, 95% *CI* [0.682, 0.765] for Network A; *r* = −0.94, *p* = 1.8 × 10^−9^, 95% *CI* [0.756, 0.845] for Network B) and lengthened decision time for correct trials (Fig. 11g, left; *r* = 0.84, *p* = 7.0 × 10^−6^, 95% *CI* [0.903, 0.963] for Network A; *r* = 0.84, *p* = 8.1 × 10^−6^, 95% *CI* [0.842, 0.931] for Network B) in both networks, while slightly shortening decision time for error trials (Fig. 11g, right; *r* = −0.81, *p* = 2.7 × 10^−5^, 95% *CI* [0.892, 0.919] for Network A; *r* = −0.89, *p* = 3.6 × 10^−7^, 95% *CI* [0.909, 0.934] for Network B). Decision confidence remained largely unchanged for both trial types in Network A, but in Network B it slightly decreased for correct trials (Fig. 11h, right; *r* = −0.73, *p* = 3.7 × 10^−4^, 95% *CI* [0.964, 0.972]) and remained constant for error trials (Fig. 11h, left).

**Fig. 11.**
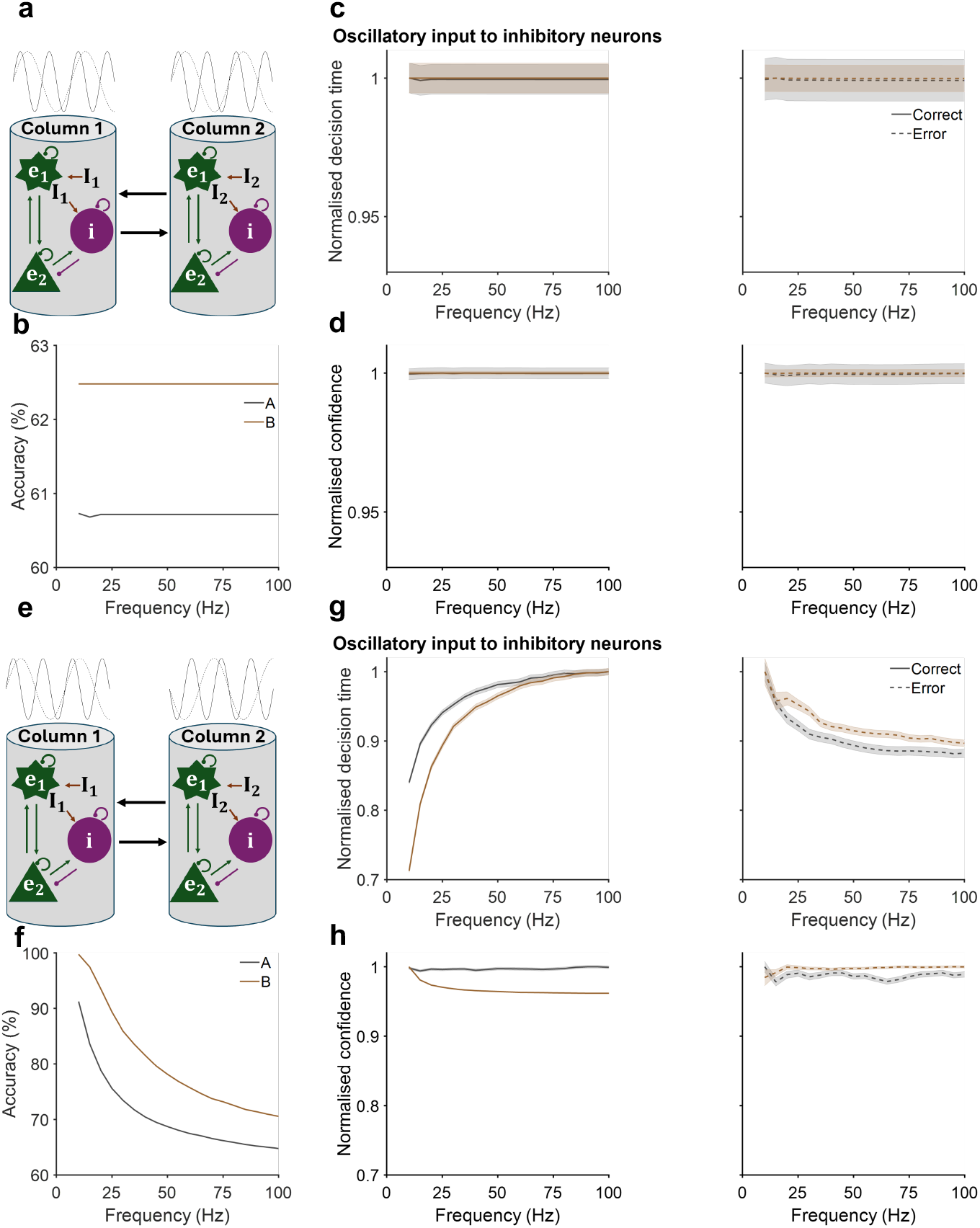
Frequency-dependent effects of in-phase and anti-phase neural oscillatory input to inhibitory neurons (amplitude = 0.08 mA) on networks with distinct time constants. **a** Schematic of in-phase sinusoidal modulation applied at varying frequencies to both cortical columns and exclusively to inhibitory neurons in each network. **b** Decision accuracy remained unchanged across frequencies in both networks. **c, d** In Network A (grey) and Network B (brown), normalised decision time (**c**) and decision confidence (**d**) remained constant for both correct (left) and error (right) trials. **e** Schematic of anti-phase sinusoidal modulation applied at varying frequencies to both cortical columns and exclusively to inhibitory neurons. **F** Decision accuracy decreased in both networks as frequency increased. **g** Normalised decision time increased for correct trials (left) and decreased for error trials (right) in both networks. **h** Normalised decision confidence remained constant for correct trials in Network A and slightly decreased in Network B; for error trials, it remained constant in both networks.

Overall, the above results (Figs. 8–11) indicate that when oscillatory input is applied to both excitatory and inhibitory neural populations, the effects of in-phase amplitude and frequency modulation primarily arise from their stronger influence on excitatory neural populations, whereas the effects of anti-phase amplitude and frequency modulation are driven mainly by their stronger influence on inhibitory neural populations during decision making.

### Network resonance enhances selective oscillatory modulation

After understanding the contribution of each neural population, we focus on the interplay between network resonance and oscillatory modulation. In particular, each network model has its intrinsic oscillatory frequency, and it is unclear whether resonance plays a role in the modulation. To quantify the potential contribution of intrinsic resonance to decision dynamics, we first computed each network’s natural frequency via (Richardson et al., 2003)

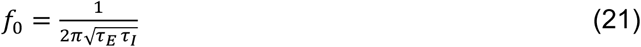

which yields *f*_0_ = 3.56 Hz for Network A and *f*_0_ = 7.12 Hz for Network B. Because our primary oscillatory drive was at 12 Hz, these baseline resonances lie outside the stimulation band, indicating that the effects observed under alpha-range inputs cannot be attributed solely to passive resonance.

To isolate resonance-specific contributions, we then re-tuned both networks so that *f*_0_ = 12 Hz and modeled their resonant response with a second-order transfer function

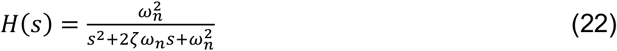

where ω_*n*_ = 2π*f*_*r*_, *ζ* is the damping ratio, and the quality factor 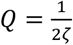 links to the bandwidth *Δf* via 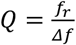. For *f* = 12 Hz and *Δf* = 6 Hz (*Q* = 2), the resonance peak has a gain *G*(*f*_*r*_) = *Q*^2^ = 4, capturing alpha-band enhancement in neural circuits (Richardson et al., 2003).

Under in-phase 12 *Hz* modulation with resonance (Fig. 12a), decision accuracy remained stable in both networks (Fig. 12b). However, Network A exhibited a markedly greater reduction in normalised decision time as amplitude increased for both types of trials (Fig. 12c; *r* = −0.92, *p* = 2.2 × 10^−9^, 95% *CI* [0.275, 0.638] for correct trials; *r* = −0.89, *p* = 7.8 × 10^−8^, 95% *CI* [0.488, 0.844] for error trials), accompanied by a stronger decline in confidence for correct and error trials (Fig. 12d; *r* = −0.94, *p* = 1.0 × 10^−10^, 95% *CI* [0.439, 0.685] for correct trials; *r* = −0.87, *p* = 2.4 × 10^−7^, 95% *CI* [0.447, 0.719] for error trials), whereas Network B still showed no significant change. During anti-phase modulation (Fig. 12e), resonance amplified the previously observed improvements: both networks attained higher accuracy at large amplitudes (Fig. 12f; *r* = 0.73, *p* = 1.8 × 10^−4^, 95% *CI* [0.902, 0.983] for Network A; *r* = 0.54, *p* = 1.0 × 10^−2^, 95% *CI* [0.933, 0.997] fo Network B), correct-trial decision time decreased (*r* = −0.93, *p* = 9.5 × 10^−10^, 95% *CI* [0.207, 0.524] for Network A; *r* = −0.91, *p* = 6.6 × 10^−9^, 95% *CI* [0.507, 0.653] for Network B) and error-trial decision time increased more steeply (Fig. 12g; *r* = 0.79, *p* = 2.0 × 10^−3^, 95% *CI* [0.737, 0.826] for Network A; *r* = 0.83, *p* = 4.0 × 10^−2^, 95% *CI* [0.939, 0.985] for Network B). Decision confidence for correct trials rose sharply in both networks (Fig. 12h, left; *r* = 0.99, *p* = 9.8 × 10^−19^, 95% *CI* [0.642, 0.801] for Network A; *r* = 0.99, *p* = 3.9 × 10^−18^, 95% *CI* [0.795, 0.870] for Network B), while error-trial confidence remained unchanged in Network B and increased in Network A (Fig. 12h, right; *r* = 0.91, *p* = 4.7 × 10^−5^, 95% *CI* [0.566, 0.753] for Network B).

**Fig. 12.**
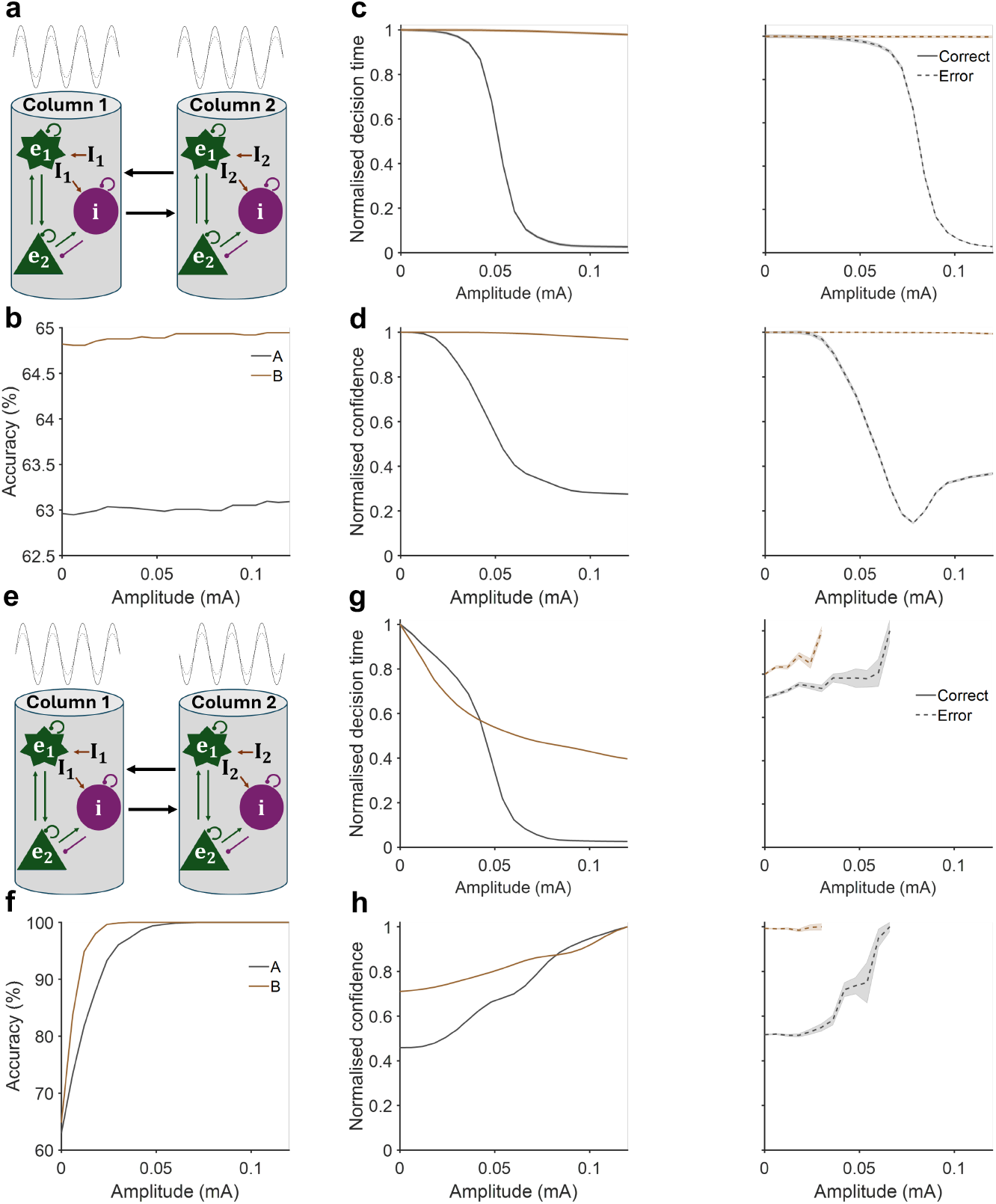
Effects of in-phase and anti-phase 12 Hz neural oscillation amplitudes on networks with differing timescales, incorporating resonance effects. **a** Illustration of in-phase sinusoidal modulation at varying amplitudes simultaneously applied to both cortical columns in each network. **b** Decision accuracy remained consistent across amplitude variations for both networks when resonance was considered. **c** With resonance, Network A (grey) exhibited a greater reduction in normalised decision time with increasing oscillation amplitude compared to conditions without resonance, consistent across both correct (left) and error (right) trials. Network B (brown) showed no significant changes in normalised decision time for either trial type with resonance included. **d** Network A demonstrated decreased normalised decision confidence with increasing oscillation amplitude, while Network B showed no significant changes in decision confidence for both correct (left) and error (right) trials. **e** Illustration of anti-phase sinusoidal modulation at varying amplitudes applied simultaneously to both cortical columns in each network, considering resonance effects. **f** Decision accuracy improved in both networks as oscillation amplitude increased. **g** Normalised decision time decreased with higher amplitude for correct trials (left) but increased for error trials (right). **h** Normalised decision confidence increased for correct trials in both networks; however, no substantial changes were observed in decision confidence for error trials in either network.

When sweeping input frequency under in-phase conditions (amplitude = 0.08 mA; Fig. 13a), accuracy remained flat across frequencies (Fig. 13b), but Network A’s normalized decision time and confidence increased sharply relative to the non-resonant case for correct and error trials within the alpha-band frequency range, and after that, it saturated at higher frequencies (Fig. 13c–d), while Network B again exhibited no sensitivity. Under anti-phase frequency modulation (Fig. 13e; *r* = −0.72, *p* = 4.4 × 10^−4^, 95% *CI* [0.655, 0.760] for Network A; *r* = −0.81, *p* = 2.9 × 10^−5^, 95% *CI* [0.687, 0.798] for Network B), both networks showed declining decision accuracy at higher frequencies (Fig. 13f). Network A’s correct-trial decision time increased more than before, while error-trial decision time decreased more sharply within the alpha-band frequency range (Fig. 13g), and both measures saturated at higher frequencies. Decision confidence in correct choices declined sharply (with a smaller decline for errors) before also saturating at higher frequencies. In contrast, Network B’s error-trial confidence remained largely flat, with only a small drop in correct-trial confidence (Fig. 13h).

**Fig. 13.**
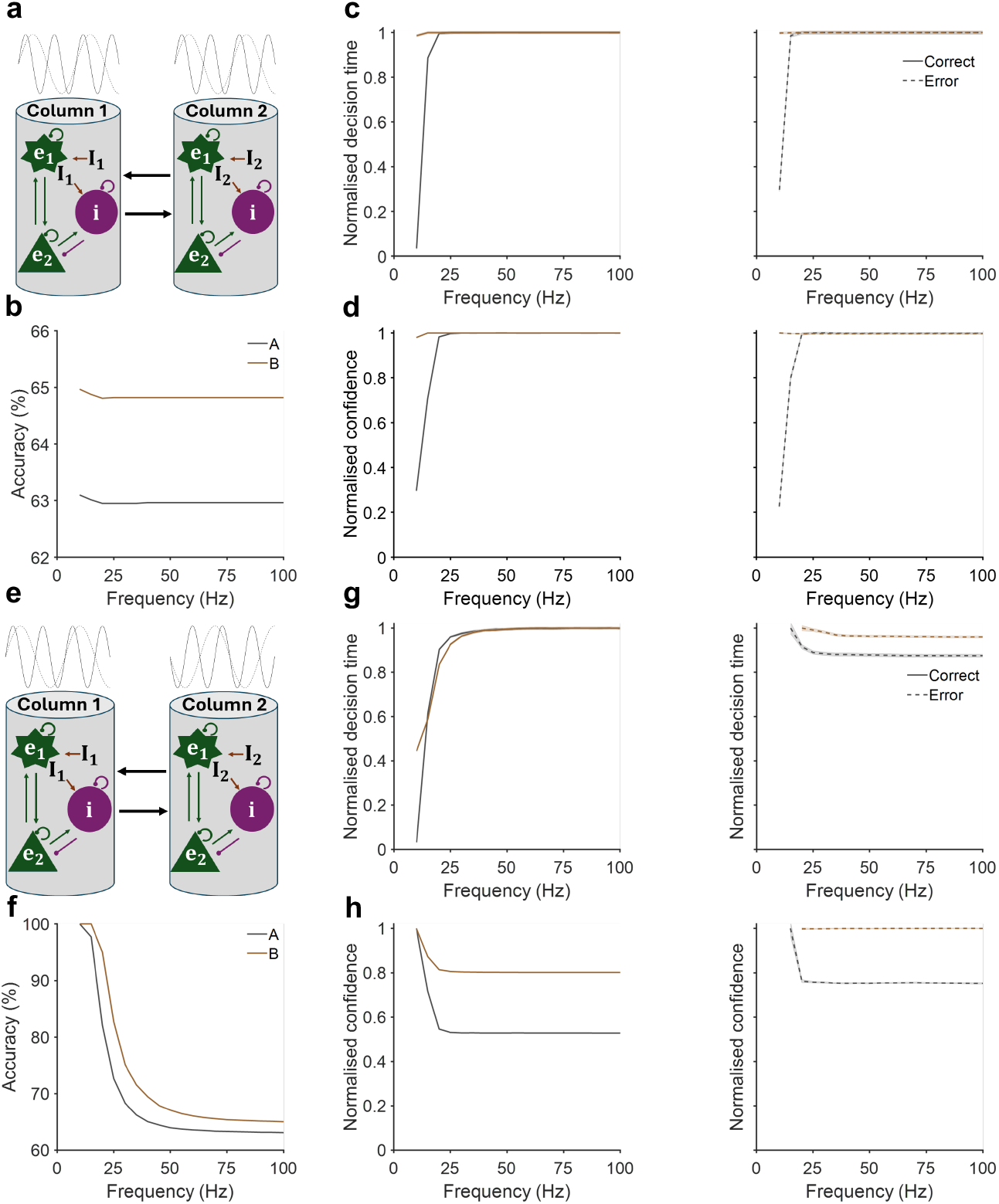
Frequency-dependent effects of in-phase and anti-phase neural oscillations (amplitude = 0.08 mA) on networks with distinct time constants, incorporating resonance effects. **a** Schematic of in-phase sinusoidal modulation applied at varying frequencies to both cortical columns in each network. **b** Decision accuracy remained unchanged across frequencies for both networks after accounting for resonance. **c, d** In Network A (grey), normalised decision time **(c)** and decision confidence **(d)** increased relative to the no-resonance model for both correct (left) and error (right) trials, whereas Network B (brown) showed no significant changes in either metric. **e** Schematic of anti-phase sinusoidal modulation at varying frequencies applied to both cortical columns, with resonance included. **f** Decision accuracy declined as oscillation frequency increased in both networks. **g** Normalised decision time increased—more than in the no-resonance model—for correct trials (left) but decreased for error trials (right). **h** Normalised decision confidence decreased for correct trials in both networks; for error trials it declined in Network A but remained largely unchanged in Network B.

Together, these results confirm that intrinsic resonance enhances the susceptibility of Network A to oscillatory inputs—particularly in the alpha band—while Network B remain relatively impermeable, even when their natural frequency is matched to the drive.

### Extension to intermediate time constants: Network C

Till now, we have only investigated the extreme physiologically allowed excitatory-to-inhibitory timescales. To explore regimes between the fast (Network A) and slow (Network B) extremes, we introduced Network C by selecting intermediate neuronal dynamical time constants (*τ*_*e*_= 20 ms; *τ*_*i*_ = 10 ms; Azimi et al., 2021) and re-tuning synaptic gains to recover bistable decision dynamics. This configuration captures a mid-range mode of cortical processing.

When subjected to in-phase and anti-phase 12 Hz drives, Network C exhibited markedly similar responses to those of Network B (Figs. 14–15). Specifically, in-phase oscillatory amplitude had negligible effects on accuracy, decision time, or confidence (Fig. 14a–d), whereas anti-phase amplitude yielded improved accuracy at higher drive strengths (*r* = 0.99, *p* = 3.1 × 10^−16^, 95% *CI* [0.726, 0.835]), with correct trials speeding up (*r* = −0.99, *p* = 4.4 × 10^−23^, 95% *CI* [0.654, 0.789]) and error trials slowing down (*r* = 0.98, *p* = 5.9 × 10^−15^, 95% *CI* [0.783, 0.870]), alongside enhanced confidence for correct choices (*r* = 0.91, *p* = 1.5 × 10^−8^, 95% *CI* [0.978, 0.985]), without altering confidence in error trials (Fig. 14e−h).

**Fig. 14.**
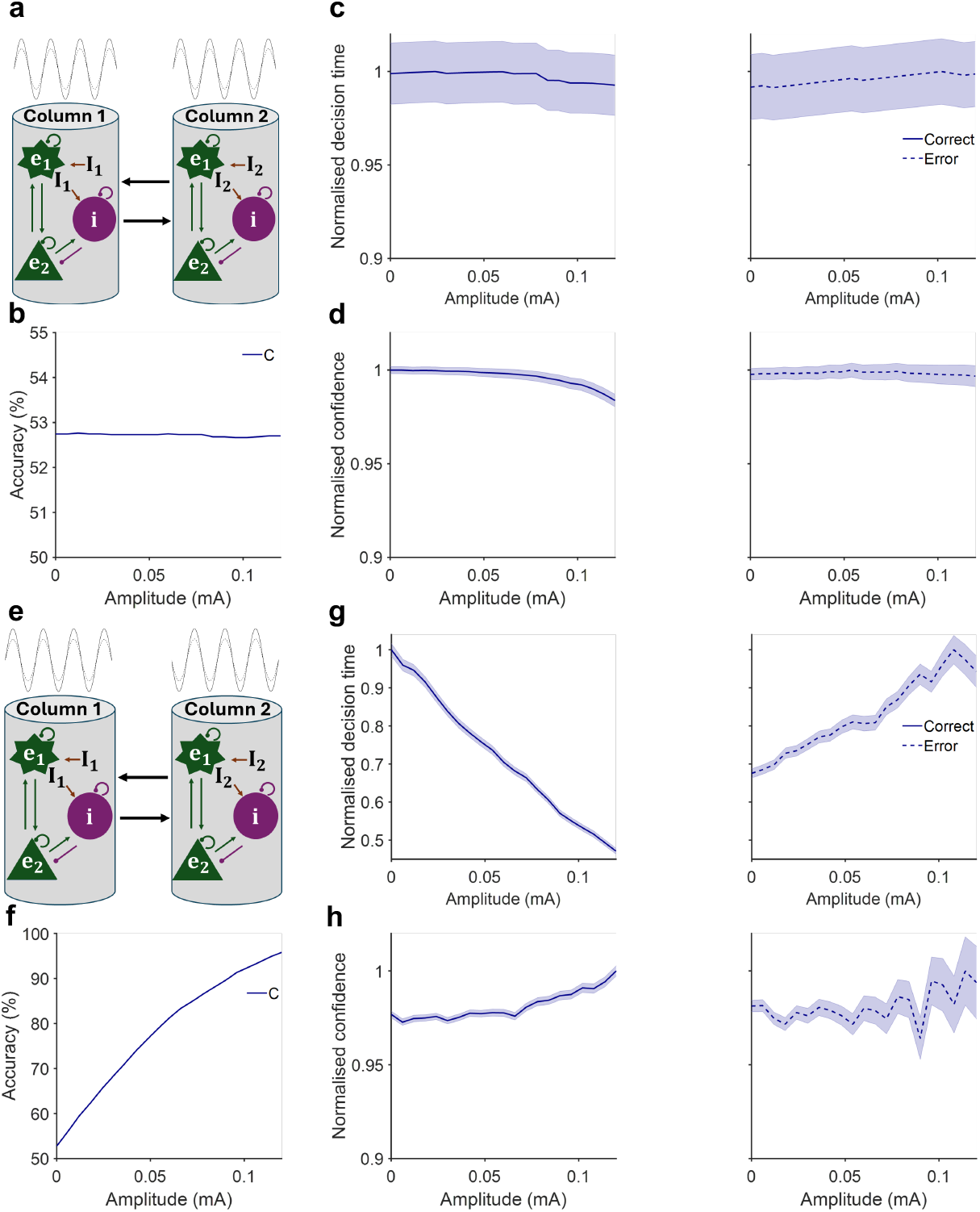
Effects of varying in-phase and anti-phase 12 Hz oscillatory amplitudes on Network C, with time constant *τ*_*E*_ = 20 ms and *τ*_*I*_ = 10 ms. **a** In-phase sinusoidal modulation at different amplitudes applied to both cortical columns for Network C. **b** Decision accuracy remained unchanged for Network C. **c** For network C (blue), showed no significant changes in normalised decision time for both trial types. **d** Network C showed no significant changes in normalised confidence for both correct (left) and error (right) trials. **e** Anti-phase sinusoidal modulation with varying amplitudes applied to both cortical columns for Network C. **f** Decision accuracy improved with higher oscillation amplitudes in Network C. **g** Normalised decision time decreased with increasing amplitude for correct trials (left), while increasing for error trials (right). **h** Normalised decision confidence increased for correct trials in Network C. However, no substantial change was observed in Network C for error trials.

**Fig. 15.**
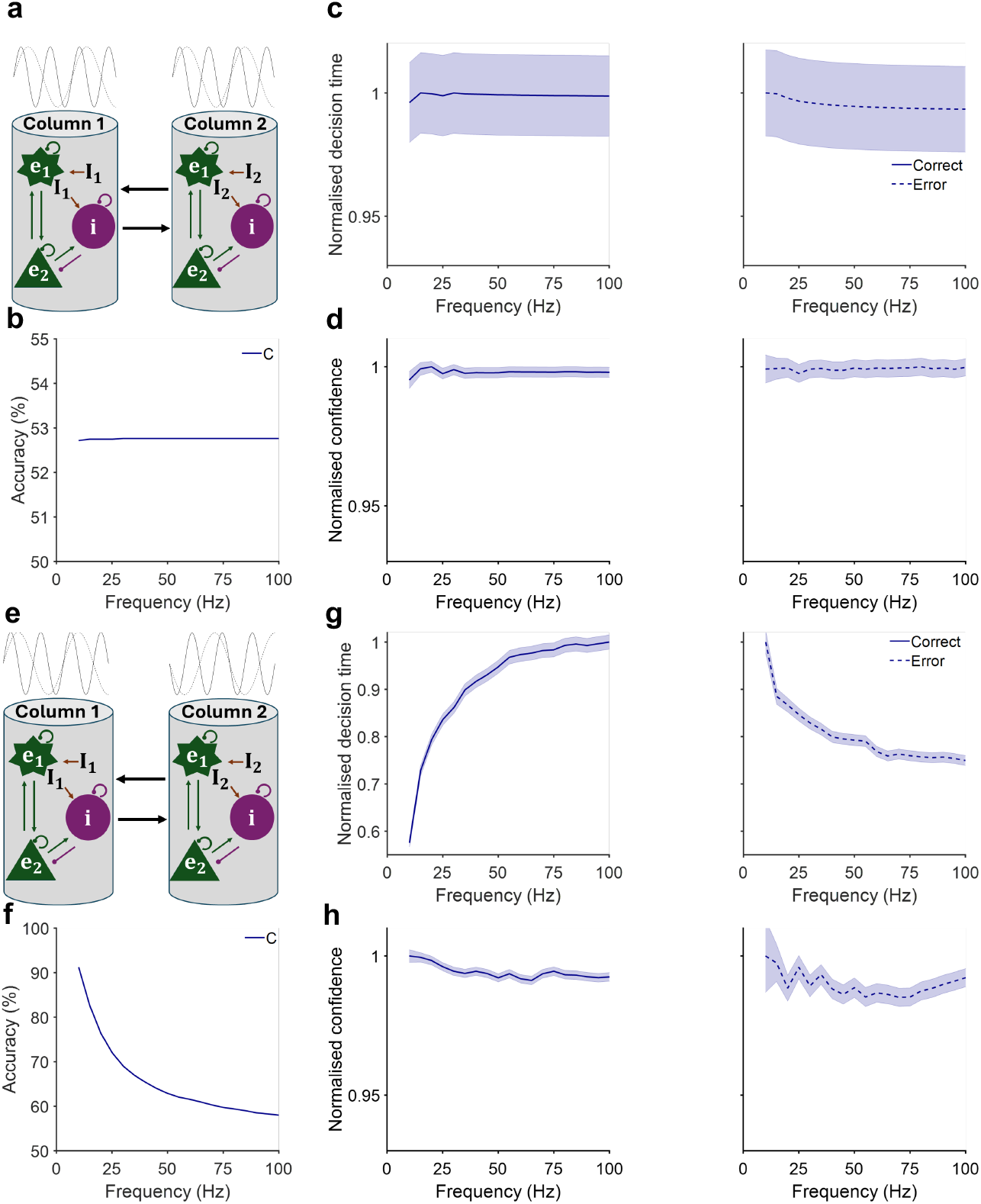
Different effects of in-phase and anti-phase neural oscillatory frequency (amplitude = 0.08 *mA*) on Network C, with time constants *τ*_*E*_ = 20 ms and *τ*_*I*_ = 10 ms. **a** In-phase sinusoidal modulation at different frequencies applied to both cortical columns for Network C. **b** Decision accuracy remained unchanged for Network C. **c, d** For Network C, neither decision time nor confidence changed significantly for either type of trials. **e** Anti-phase sinusoidal modulation with varying frequencies applied to both cortical columns for Network C. **f** Decision accuracy declined with higher oscillation frequencies in Network C. **g** Normalised decision time increased with increasing amplitude for correct trials (left), while it decreased for error trials (right). **H** Normalised decision confidence slightly decreased for correct trials in Network C, and showed no substantial change for error trials.

Likewise, in-phase frequency modulation (0.08 mA amplitude) did not alter performance (Fig. 15a–d), but anti-phase frequency increases degraded accuracy (*r* = −0.87, *p* = 1.6 × 10^−6^, 95% *CI* [0.621, 0.720]), prolonged correct-trial decision times (*r* = 0.85, *p* = 5.0 × 10^−6^, 95% *CI* [0.769, 0.911]), shortened error decision times (*r* = −0.85, *p* = 4.0 × 10^−6^, 95% *CI* [0.778, 0.834]), and modestly reduced correct-trial confidence (*r* = 0.90, *p* = 1.8 × 10^−7^, 95% *CI* [0.993, 0.995]) without affecting errors (Fig. 15e−h).

These results indicate that Network C mirrors Network B’s susceptibility to anti-phase drives. Crucially, the differential oscillatory sensitivity across Networks A–C suggests that empirical measurements of phase- and amplitude-dependent modulation *in vivo* could serve as a “synaptic fingerprint”, revealing whether a given cortical area operates in a fast, relay-like mode or in a slower, integrative regime during decision-making.

### Neural circuit mechanisms of oscillation-induced decision modulations

Next, we seek to understand the network dynamical mechanisms underlying the above simulated results. Given that phase differences generally affect decision behaviours in a progressive manner, it suffices to provide a conceptual understanding for the two extreme cases: in-phase and anti-phase neural oscillatory modulations. This can be investigated by projecting the trial-averaged trajectories onto lower (two) dimensional state (decision) space. For simplicity, we focused only with stimulus bias of 7%.

Our simulations generally indicated that, for network A, in-phase modulation of neural oscillation with higher amplitudes tended to retain the network state along the decision uncertainty manifold (i.e., along the state-space diagonal) in state space (Fig. 16a), hence, decreasing decision confidence. Concurrently, larger oscillation amplitudes led to larger deflection in phase space, leading to greater momentum towards the choice attractors without favouring any of the choices. Therefore, these explain why for network A, in-phase amplitude-modulation decreased decision confidence while expediting decision formation, without influencing decision accuracy (Fig. 2).

**Fig. 16.**
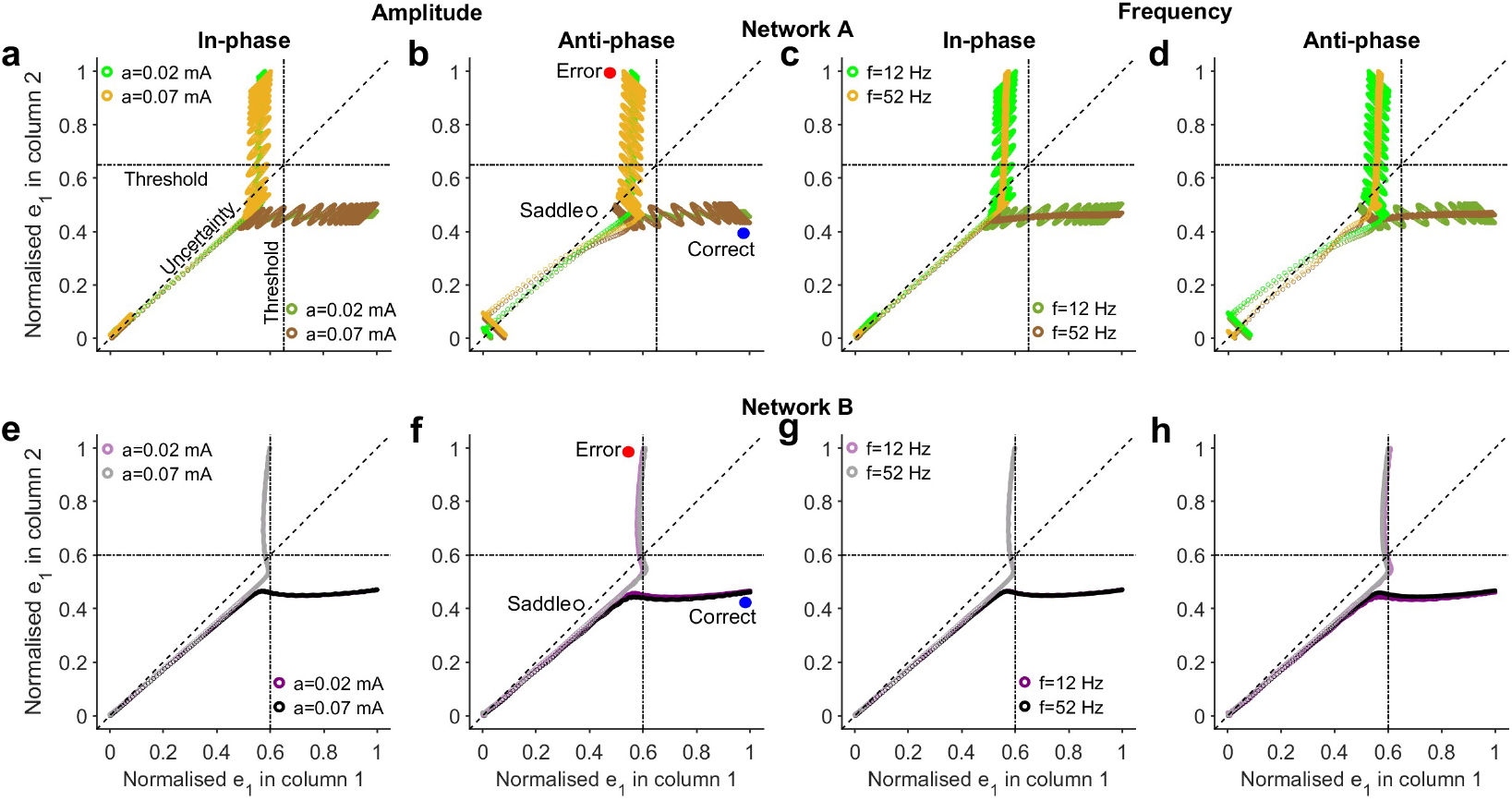
Neural oscillations on choice behaviour in (projected) state space. In state (EPSP) space, the trajectories denote neural population EPSPs (normalised *e*_1_ in column 1 and normalised *e*_1_ in column 2), averaged across correct or error trials (with 7% stimulus bias). Top: Network A. Higher amplitude/frequency, brown (gold) trajectories indicate correct (error) trials; lower amplitude/frequency, dark (light) green for correct (error) trials. Bottom: Network B. Higher amplitude/frequency, dark (light) grey trajectories indicate correct (error) trials; lower amplitude/frequency, dark (light) purple for correct (error) trials. **a-d** Modulation due to in-phase (**a, c**) and anti-phase (**b, d**) with different oscillation amplitudes. **e-h** Modulation due to in-phase (**e, g**) and anti-phase (**f, h**) with different oscillation frequencies. Filled (unfilled) circle: stable choice attractor (metastable saddle-like steady state); blue (red) for correct (error) choice. Although indicated only in (**b**) and (**d**), they exist, in the same positions, in other panels (not indicated for clarity). Time spacing between consecutive trajectory points for network A (B) is 1 ms (5 ms).

Conversely, anti-phase neural oscillation with higher amplitudes provided swinging momentum towards the choice attractors for network A, with a greater tendency for correct choices (Fig. 16b), and leading to more accurate decisions. Further, higher oscillatory amplitude exerted a greater tendency to move the network state (orthogonally) away from the decision uncertainty manifold, increasing decision confidence. Although one might consider anti-phase modulation with higher oscillation amplitude to also deflect trajectories more in state space, for error choices, the network state also passed closer towards a metastable saddle steady state (Fig. 16b), which in turn slowed trajectories (Wong and Wang, 2006; Wong et al., 2007) and leading to slower error decision times. Therefore, these explain why for network A, anti-phase amplitude-modulation increased decision accuracy and confidence, and expedited decision formation for correct trials while slowing error trials (Fig. 4).

For network B, in-phase oscillatory modulatory effects were less prominent with higher oscillation amplitude (Fig. 16c) due to the slow excitatory synapses filtering out oscillatory inputs, making the network more robust to such perturbations (Fig. 2). However, with anti-phase oscillatory modulation, larger oscillation amplitude increased the tendency to slightly move towards the correct attractor faster but slower for the error attractor, while also moving away from the uncertainty manifold (Fig. 16d). Thus, for network B, anti-phase modulation with increasing amplitude increased decision accuracy and confidence while expediting decision formation for correct trials and slowing error trials (Fig. 4).

In terms of in-phase oscillation with varying frequency, network A’s state tended to remain along the decision uncertainty manifold in state space at lower frequencies and slower towards choice attractors without biasing towards a specific attractor (Fig. 16e). Therefore, with network A, in-phase modulation with increasing frequency increased decision confidence while slowing decision formation, without influencing decision accuracy for network A (Fig. 3).

In contrast, albeit non-intuitively, anti-phase neural oscillation with lower frequency manifested as if they had larger trial-averaged trajectory fluctuations akin to larger oscillation amplitudes (compared with Figs. 7a-b). Hence, the explanations of the simulated behaviour were reverse of that with higher oscillation amplitude (see above).

Thus, for network A, anti-phase oscillatory modulation with increased frequency decreased decision accuracy and confidence while increasing decision time (Fig. 5). For network B, in-phase oscillatory modulation was not noticeable with increasing frequency (Fig. 16g), explaining the simulated results in Fig. 3. However, for anti-phase oscillatory modulation with increasing oscillation frequency, it had the opposite, albeit subtle, effect as with increasing oscillation amplitude (compare Fig. 16h with Fig. 16d). This explained the reverse trends observed in Fig. 5.

## Discussion

In this study, we developed a cortical-column decision network with multiple interacting populations and self-feedback to probe how ongoing neural oscillations shape perceptual decisions. In particular, we have developed the first functional decision-making network (Network A) model with slower inhibitory than excitatory synapses. By contrasting it with another network (Network B) with slower excitatory than inhibitory synapses, we identified distinct intrinsic and emergent timescales that differentially modulate the influence of oscillations on decision accuracy, speed, and confidence (Figs. 1–7; Figs. S1–S2). Across systematic manipulations of phase, amplitude, and frequency, oscillations selectively dissociated these behavioural measures and, in specific operating regimes, violated the speed–accuracy trade-off (SAT) (Figs. 2–6) (Bogacz et al., 2010; Heitz, 2014). These effects depended on synaptic composition and emergent neural circuit dynamics.

Consistent with human EEG work, in-phase 12 Hz (alpha) modulation reduced decision confidence without altering accuracy in Network A (Fig. 2d), with the strongest effects for the lower frequency bands (Fig. 3d; Fig. 6b). This pattern supports a role for alpha—and low frequencies more generally—in regulating the flow of sensory information during choice (Samaha et al., 2017, 2020), potentially via phasic gating of evidence integration. Importantly, in-phase oscillations above ∼30 *Hz* exerted less influence, consistent with effective high-pass filtering by synaptic dynamics; likewise, in anti-phase conditions, oscillatory effects waned beyond ∼75 *Hz* (Figs. 5-6). Differential entrainment analyses further clarified why Network A is more susceptible than Network B: under identical 12 *Hz* drive, Network A showed strong phase-locked responses whereas Network B displayed weak, shifted, and intermittent coupling (Fig. S2), reinforcing the concept that intrinsic timescales constrain how rhythms access neural (decision) computations (Buzsáki, 2006; Fries, 2015).

Physiologically, anti-phase relationships across nearby subcircuits could arise from travelling waves from one brain location to another, signal conduction delays interacting with heterogeneous time constants, push–pull lateral inhibition, or laminar phase reversals (Rubino et al., 2006; Maier et al., 2010; Li and Zhou, 2011). In this work, anti-phase modulation produced a distinct profile: increasing anti-phase amplitude improved decision accuracy and confidence and speed of correct choices while slowing error choices (Fig. 4b–d), with broadly similar trends in both networks but larger effects in Network A. Frequency sweeps revealed the complementary pattern: increasing anti-phase frequency progressively degraded decision accuracy and confidence and slowed correct trials (Fig. 5b–d), mirroring the amplitude–frequency opponency seen throughout the maps in Figs. 6 (theta– low gamma band dominance in Network A and broad theta– high gamma accuracy benefits in Network B).

Intriguingly, these anti-phase benefits parallel neurostimulation results in which anti-phase alpha transcranial alternating current stimulation (tACS) across early visual areas improved motion discrimination compared with in-phase or sham, suggesting that temporal offsets between brain regions can reduce mutual interference and sharpen evidence integration (Salamanca-Giron et al., 2021). More generally, our modelling results are consistent with experimental evidence demonstrating the role of alpha oscillations in modulating perceptual sensitivity (Samaha et al., 2017, 2020) and the roles of beta/gamma oscillations in evidence encoding and action planning (Donner et al., 2009; Siegel et al., 2011). By varying the inter-column phase difference (Δ*φ*), it revealed graded and selective control of behaviour. When the column encoding the correct choice led in phase, accuracy rose and decision time fell with increasing Δ*φ* and amplitude, whereas error decision time either increased or remained stable (Fig. S3). The converse manipulation (error column leading) reversed these trends (Fig. S4). Frequency-by-phase analyses reinforced these relationships (Figs. S5−S6), and stimulus-phase alignments showed that sensory timing relative to ongoing inter-column phase further gates outcomes (Fig. 7).

These mechanisms provide concrete routes by which phase-lead identity and timescale diversity implement the selective acceleration of correct trials and the selective slowing of errors that we observe (Figs. 4, 6; Figs. S3–S6). Our phase-dependent stimulus timing results (Fig. 7) suggest the possibility of experimental tests using controlled sensory phase and inter-columnar phase lags (Mathewson et al., 2009; Strauß et al., 2015). Behaviourally, such patterns constitute principled SAT violations—simultaneous gains in speed and accuracy when the correct column leads—akin to contexts where initial choice bias or high-stakes contingencies decouple speed and accuracy (Busemeyer, 1993; Shevlin et al., 2022). Mechanistically, these results formalise oscillatory phase structure as a physiological substrate for SAT reconfiguration.

Our work was guided by known synaptic timescales governed by GABA-, NMDA- and AMPA-receptors, while acknowledging that purely GABA_B_-mediated synapses are rare and mixed GABA_A_/GABA_B_ motifs are common (Tamás et al., 2003; Oláh et al., 2009), and that classical studies already emphasised diverse inhibitory microcircuits (Connors et al., 1988). Although we have explored the two physiologically plausible extreme cases of excitatory vs inhibitory time constants, we have also investigated intermediate values and similar results showed.

To elucidate neuronal mechanisms, we implemented inputs exclusively to each neural population (Figs. 8–11). For in-phase modulation, the effects were mediated predominantly via excitatory neural populations in Network A, with reduced decision confidence and faster correct choices. In contrast, for anti-phase modulation, the effects were contributed mainly by inhibitory neural populations, with decision accuracy gains, faster correct choices and slower error choices in both networks. These findings support the idea that inhibitory timing governs synchrony and routing, while excitatory channels set integrative timescale (Buzsáki and Wang, 2012; Soltani et al., 2021). These results could perhaps be validated in *in vivo* experiments using electrical microstimulation or optogenetics approach.

To shed light on the network dynamical mechanisms, we incorporated intrinsic resonance and found that alpha-band susceptibility to be amplified predominantly in Network A, sharpening both in-phase and anti-phase signatures while leaving slow circuits comparatively robust, even when their natural frequency was matched to the drive (Figs. 12–13).

To align with physiological neuronal membrane time constants (Bal and Destexhe, 2009), we developed another network model (Network C) with excitatory and inhibitory time constants of *τ*_*e*_= 20 ms and *τ*_*i*_ = 10 ms, respectively (Azimi et al., 2021), which sit midway between the previously studied fast and slow excitatory-to-inhibitory time constants (Network A and B). Such smaller time constants have been shown to exhibit decision dynamics (Roxin and Ledberg, 2008), but requires finer tuning of parameters close to bifurcation point (Wong and Wang, 2006). After tuning the synaptic gains to exhibit bistable decision dynamics, we found that Network C mirrors Network B’s susceptibility to anti-phase drives (Figs. 14-15). Hence, these indicate that our modelling approach was chosen to capture the functional distinction between fast and slow inhibitory dynamics, and represents a computational abstraction, agnostic to specific synaptic types.

We further probe for neural circuit mechanisms using state-space analyses (Fig. 16) to explain various observed decision modulatory effects. We found in-phase modulation tends to keep Network A’s trajectories near the decision-uncertainty manifold (state-space diagonal), reducing the instantaneous separation between choice states and thereby lowering confidence while modestly hastening choices via added momentum (Fig. 16a). In contrast, anti-phase modulation swings trajectories away from the uncertainty manifold toward the choice attractors, particularly more towards the correct choice attractor (with larger and deeper basin of attraction (Wong and Wang, 2006)), jointly increasing accuracy and confidence and speeding correct choices (Fig. 16b).

For error trials under anti-phase input, paths pass closer to metastable saddle regions, selectively delaying errors (Fig. 16b) (see also Wong and Wang, 2006 and Wong et al., 2007). These dynamical signatures complement diffusion-type accounts by exposing how rhythmic push–pull forces sculpt vector fields and path geometry during bounded evidence accumulation (Wong and Wang, 2006; Ratcliff and McKoon, 2008). The weaker in-phase effects in Network B (Fig. 16c) and its modest anti-phase biasing (Fig. 16d) follow from slower excitatory dynamics that filter rapid rhythmic perturbations. Frequency manipulations invert these amplitude-driven patterns (Fig. 16e–h), consistent with the opponency observed behaviourally (Figs. 3, 5 and 6). Together, these provided deep conceptual understanding for the observed phenomena in both our model simulations and experimental studies (Donner et al., 2009; Samaha et al., 2017, 2020; Salamanca-Giron et al., 2021).

Our study highlights several directions for future research. We have modelled oscillations as sinusoids and did not include explicit cross-frequency interactions or irregular dynamics that characterise more realistic cortical rhythms (Wang, 2010; Voytek et al., 2015). We also did not model the generators of the rhythms themselves or with long-range delayed coupling between columns (Jansen and Rit, 1995; Deco et al., 2009). Extending the framework to state- and context-dependent neuromodulation and disorder-specific circuit changes should further bridge computational predictions with systems and clinical neuroscience.

In sum, oscillatory phase, amplitude, and frequency emerge as tuneable control parameters that can decouple—and even invert—relationships among speed, accuracy, and confidence. These effects selectively depend on synaptic timescales and on which neural population leads in phase, providing a mechanistic route by which ubiquitous brain rhythms flexibly reconfigure decision computations (Buzsáki, 2006; Fries, 2015). By mapping these dependencies and linking them to state-space geometry, our results motivate targeted neuromodulation strategies and provide testable predictions for how rhythms sculpt choice in health and disease.

## Acknowledgements

A.A. and K.W.-L. were supported by HSC R&D (STL/5540/19) and MRC (MC_PC_20020).

