## Supplementary Information for "Neural oscillation as a selective modulatory mechanism on decision confidence, speed and accuracy"

**Supplementary Information**

This supplementary file provides extended methodological details, model schematics, and additional simulation results that complement and support the findings presented in the main manuscript. The supplementary figures collectively illustrate how neural oscillatory modulation—applied with varying amplitudes, frequencies, and phase relationships—affects decision-making behaviours such as accuracy, decision time, and confidence within a two-column cortical neural mass model. Results are presented for networks with different intrinsic timescales (Networks A, B, and C), with separate analyses for oscillatory inputs targeting excitatory or inhibitory populations. Several figures explore phase-lead and phase-lag conditions between cortical columns, as well as frequency-specific effects spanning theta to alpha ranges. Additional simulations also incorporate resonance effects to assess their influence on oscillatory modulation outcomes. Together, these supplementary materials provide a comprehensive view of the model's dynamic responses across diverse modulation regimes.

**Fig. S1**: Detailed schematic of the two-column cortical neural mass model with symmetrical connectivity, plus EPSP comparison for Networks A and B across evidence qualities.

**Fig. S2**: Distinct modulation patterns in Networks A and B under 12 Hz oscillatory input to excitatory and inhibitory neurons, showing differences in entrainment, spectral peaks, and coupling strength.

**Fig. S3**: Phase-difference effects (12 Hz, column 1 leading) across amplitudes on decision accuracy, time, and confidence in Networks A and B.

**Fig. S4**: Phase-difference effects (12 Hz, column 2 leading) across amplitudes on decision accuracy, time, and confidence in Networks A and B.

**Fig. S5**: Phase-difference effects across frequencies (column 1 leading) on decision measures in Networks A and B.

**Fig. S6**: Phase-difference effects across frequencies (column 2 leading) on decision measures in Networks A and B.

**a**

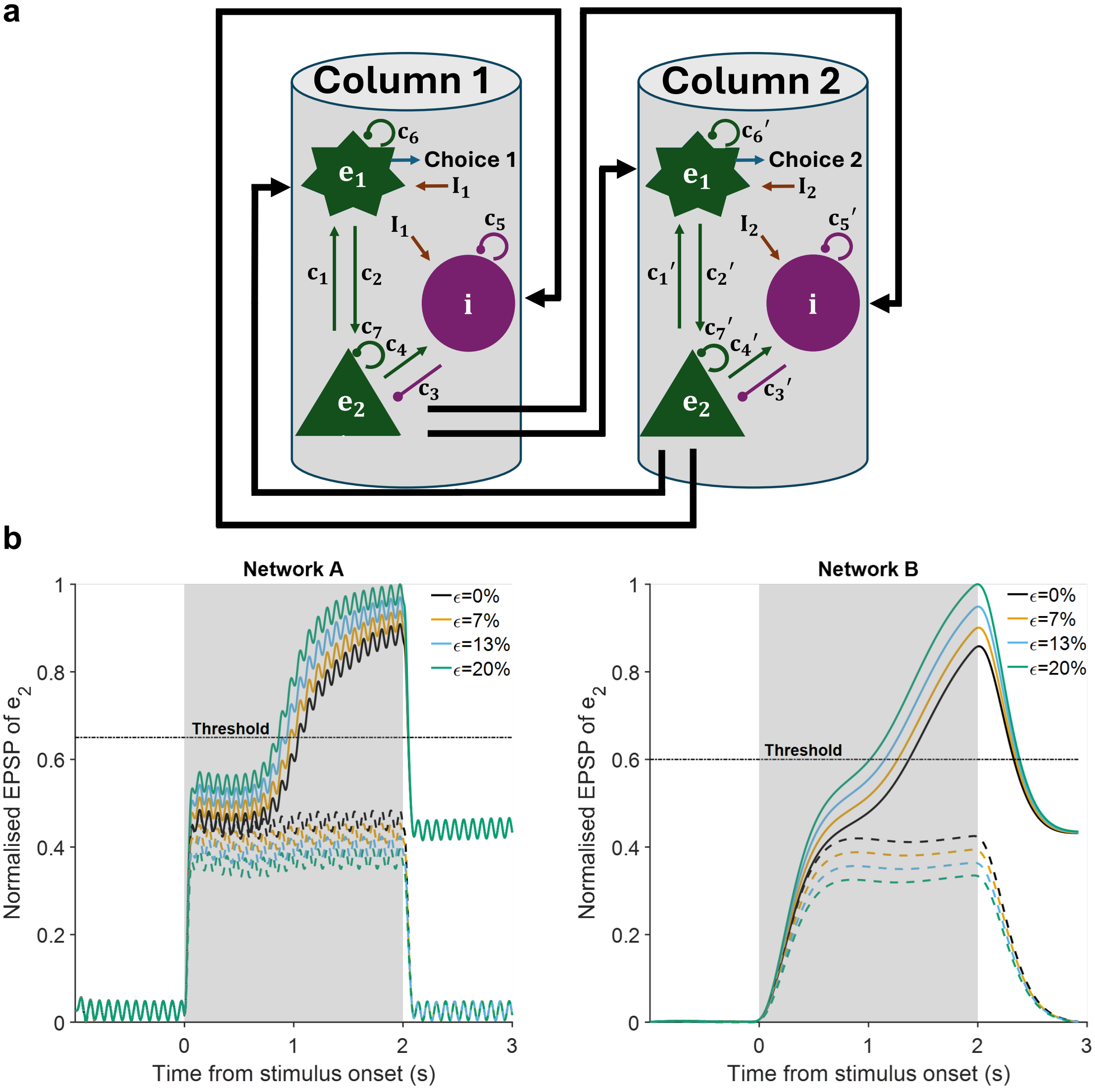


**Fig. S1** A detailed schematic diagram of a two-column cortical neural mass model for two-choice decision-making. **a** In this model, each column contains two excitatory populations ($e_1$ and $e_2$) along with a single inhibitory population (i). Inputs $I_1$ and $I_2$ are provided to both the excitatory and inhibitory neural populations, and lateral interactions are incorporated across the columns. c's denote connectivity strengths, including within-column connectivity and self-connectivity. No intrinsic bias between columns: cross-column connectivity is symmetrical, and within-column connectivity in one column mirrors that of the other column. **b** Normalised EPSP of neuron $e_2$ averaged over correct trials for different evidence qualities $\epsilon$ in Network A (left) and Network B (right).

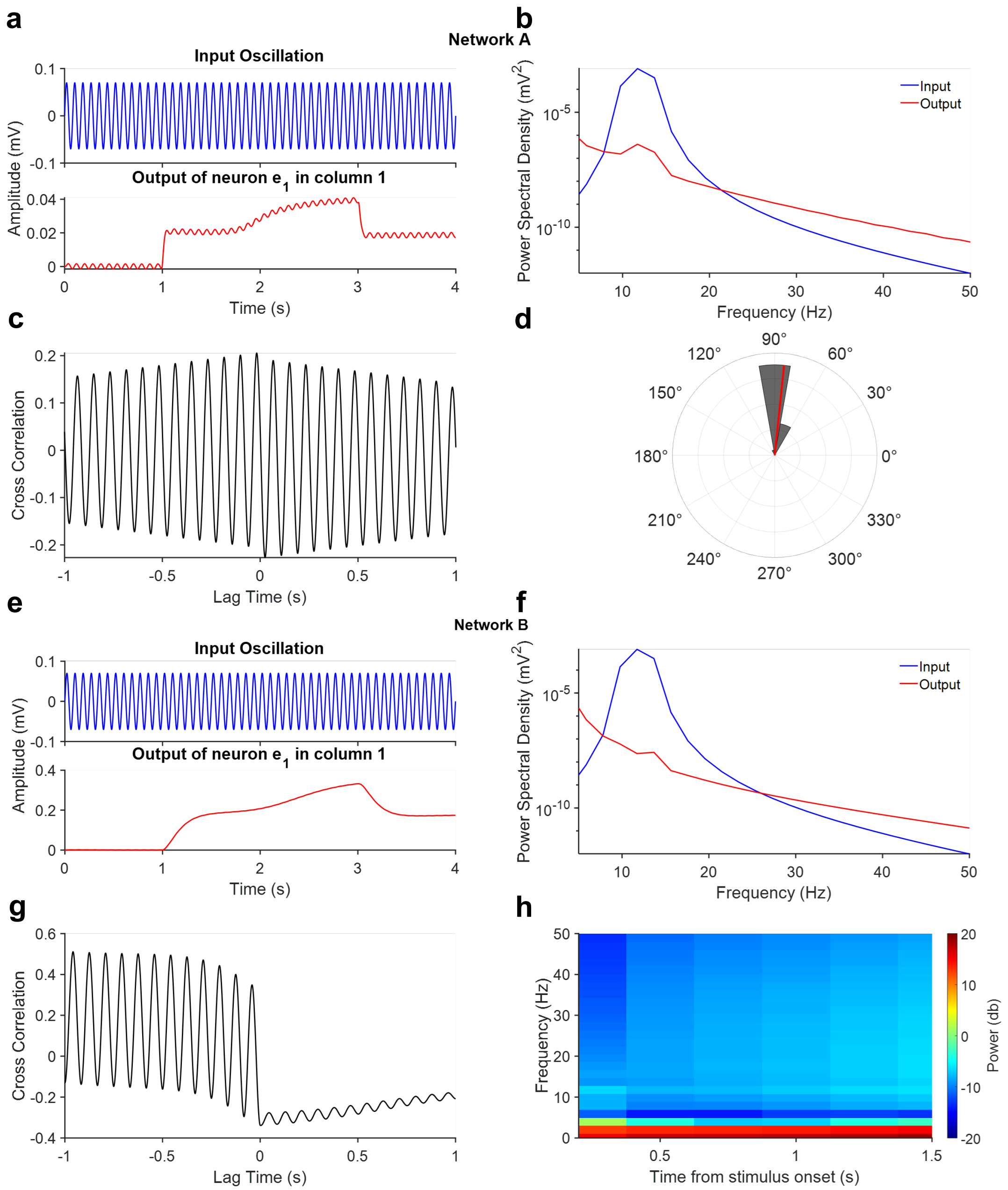


**Fig. S2** Oscillatory input modulates Networks A and B in distinct ways. **a** Network A – *top:* oscillatory drive (12 Hz) applied to excitatory neuron $e_1$ and the inhibitory population $i$; bottom*:* output of $e_1$ in column 1, showing pronounced entrainment by the input. **b** Power-spectral density (PSD) of the signals in **a**, with a clear alpha-band peak (~12 Hz) in both the input and the $e_1$ output. **c** Cross-correlation between input and $e_1$ output in Network A demonstrates strong coupling; the output lags the input by ~25 ms. **d** Phase–locking analysis confirms robust modulation in Network A (mean resultant vector length = 0.9927) at a preferred phase of 84.1°. **e** Network B – top*:* the same 12 Hz drive applied to $e_1$ and $i$; bottom*:* $e_1$ output in column 1, which is only weakly modulated. **f** Power spectral density for *Network B:* the input retains its ~12 Hz alpha peak, whereas the $e_1$ output peak shifts to ~14 Hz. **g** Cross-correlation in *Network B* reveals much weaker input-output coupling. **h** Spectrogram of the $e_1$ output in column 1 *(*Network B*)* shows only faint alpha-band modulation.

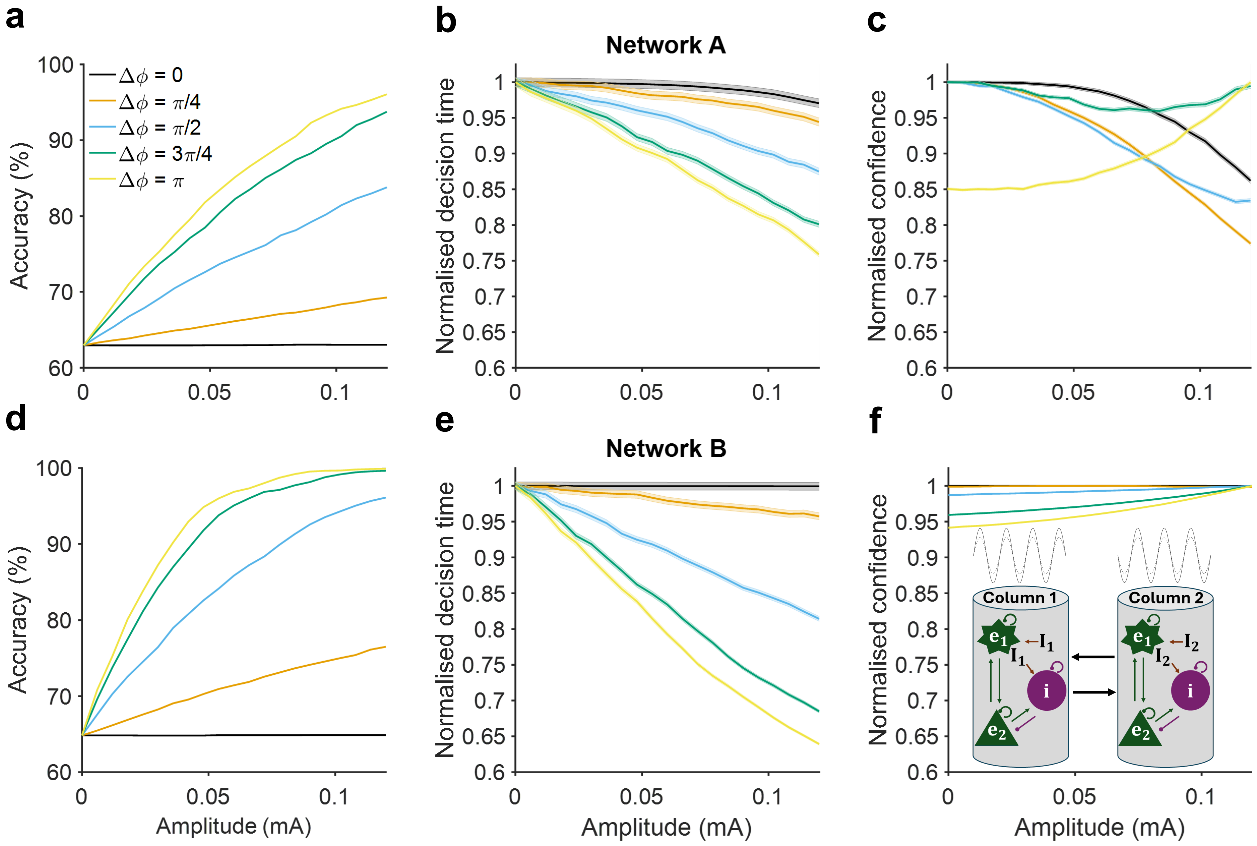


**Fig. S3** Effects of oscillatory modulation on decision behaviours across amplitudes and phase differences at 12 Hz (alpha band), with oscillatory input to column 1 (correct decision) leading that of column 2 (error decision). **a-c** Effects on network A. As phase difference between oscillatory inputs to columns increased, decision accuracy progressively increased with oscillation amplitude (**a**)**,** decision time progressively decreased with oscillation amplitude (**b**), and decision confidence changed from decreasing to increasing trends with oscillation amplitude (**c**). **d-f** Effects on network B. Same effects on decision accuracy (**d**) and decision time (**e**) as network A, but decision confidence only increased with oscillation amplitude (**f**). Inset: Schematic of type of oscillatory modulation.

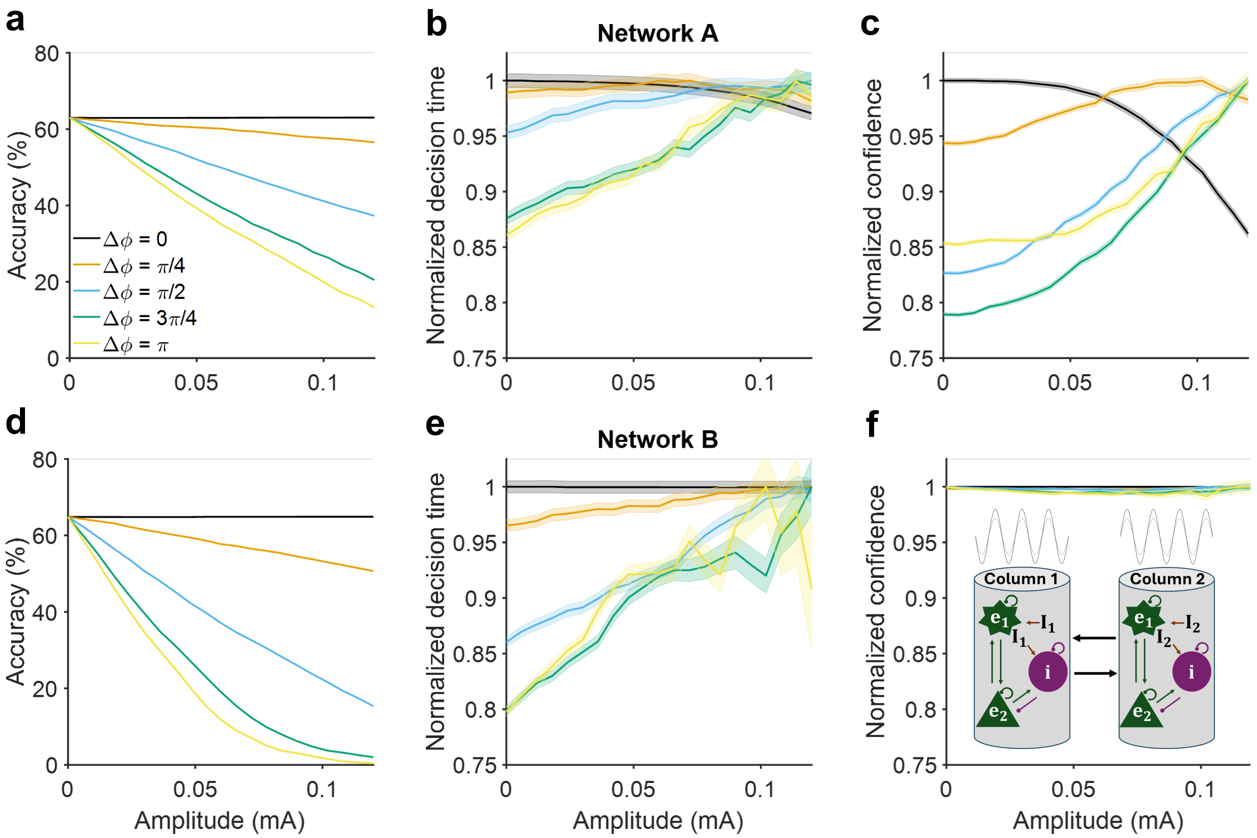


**Fig. S4** Effects of oscillatory modulation on choice behaviours across amplitudes and phase differences at 12 Hz (alpha band), with oscillatory input to column 2 (error decision) leading that of column 1 (correct decision). **a-c** Effects on network A. As phase difference between oscillatory inputs to columns increased, decision accuracy progressively decreased with oscillation amplitude (**a**), while decision time (**b**) and decision confidence (**c**) changed from decreasing to increasing trends with oscillation amplitude. **d-f** Effects on network B. Same effect on decision accuracy as network A (**d**), but decision time only increased with oscillation amplitude (**e**) and decision confidence remained constant (**f**). Inset: Schematic of type of oscillatory modulation.

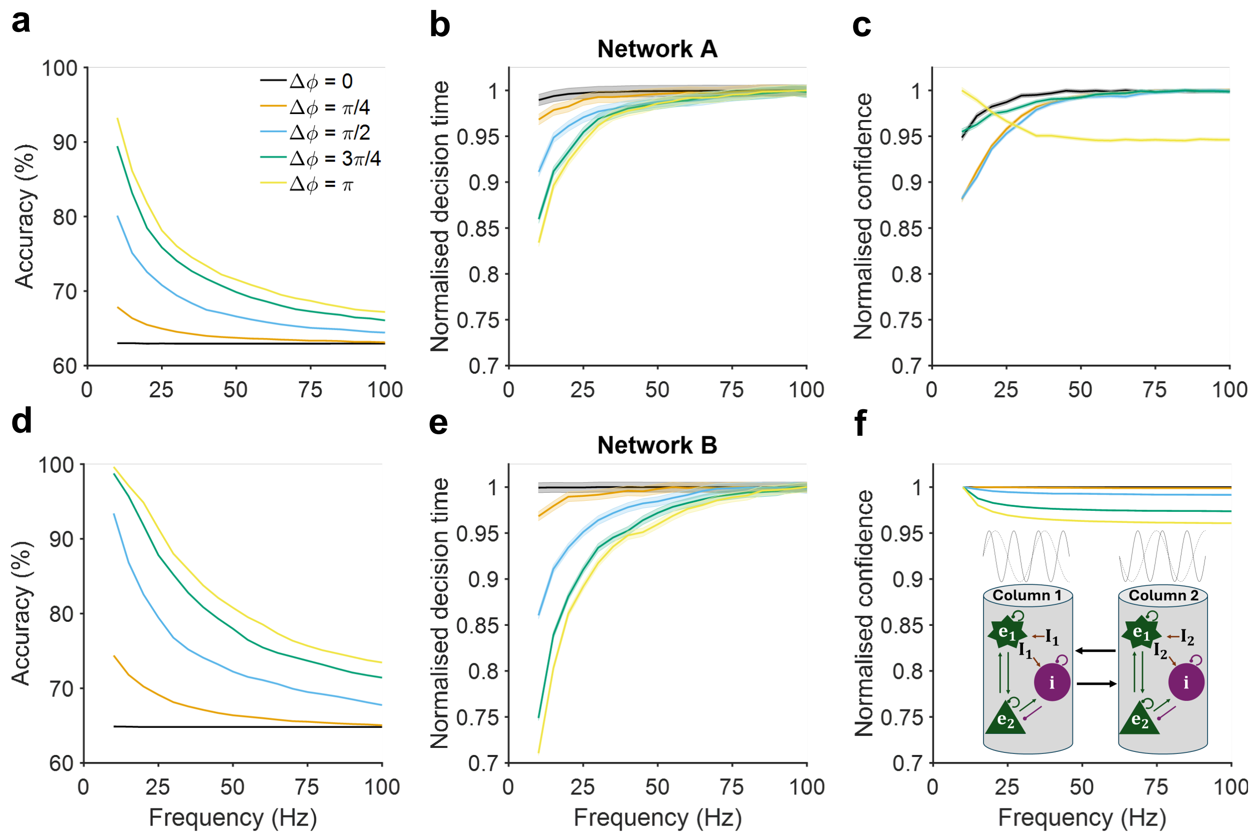


**Fig. S5** Effects of oscillatory modulation on decision behaviours across frequencies and phase differences, with oscillatory input to column 1 (correct decision) leading that of column 2 (error decision). **a-c** Effects on network A. As phase difference between oscillatory inputs to columns increased, decision accuracy progressively increased but with decreasing trend with oscillation frequency (**a**), and decision time decreased but with increasing trend with oscillation frequency (**b**), while decision confidence changed from increasing to decreasing trends with oscillation frequency (**c**). **d-f** Effects on network B. Same effects on decision accuracy (**d**) and decision time (**e**) as network A, but decision confidence only slightly decreased with oscillation frequency (**f**). Inset: Schematic of type of oscillatory modulation.

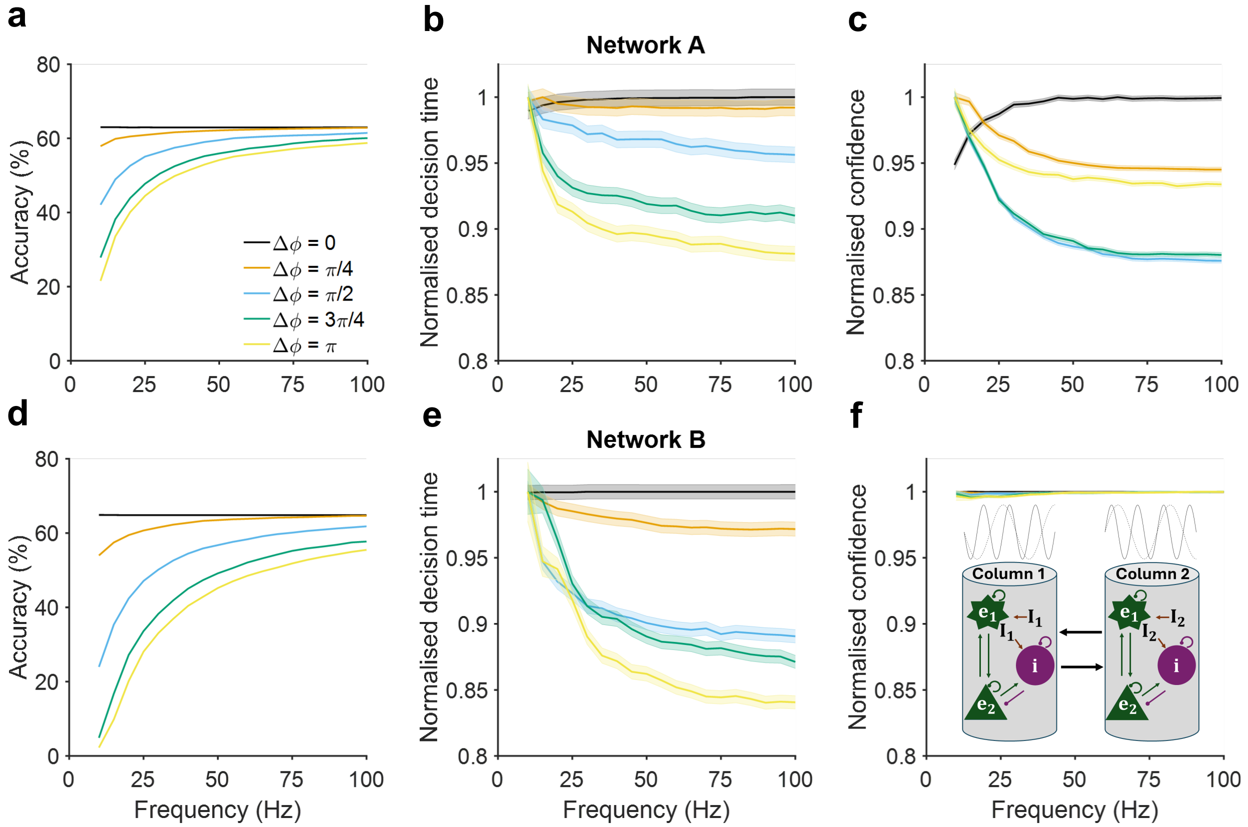


**Fig. S6** Effects of oscillatory modulation on decision behaviours across frequencies and phase differences, with oscillatory input to column 2 (error decision) leading that of column 1 (correct decision). **a-c** Effects on network A. As phase difference between oscillatory inputs to columns increased, decision accuracy progressively decreased but with increasing trend with oscillation frequency (**a**), decision time decreased with oscillation frequency (**b**), while decision confidence changed from increasing to decreasing trends with oscillation frequency (**c**). **d-f** Effects on network B. Same effects on decision accuracy (**d**) and decision time (**e**) as network A, but decision confidence remained constant (**f**). Inset: Schematic of type of oscillatory modulation.
